# A systematic comparison of current bioinformatic tools for glycoproteomics data

**DOI:** 10.1101/2022.03.15.484528

**Authors:** Valentina Rangel-Angarita, Keira E. Mahoney, Deniz Ince, Stacy A. Malaker

## Abstract

Glycosylation is one of the most common post-translational modifications and generates an enormous amount of proteomic diversity; changes in glycosylation are associated with nearly all disease states. Intact glycoproteomics seeks to determine the site-localization and composition of glycans along a protein backbone via mass spectrometry. Following data acquisition, raw files are analyzed using search algorithms to define peptide sequence, glycan composition, and site localization. Glycoproteomics is rapidly expanding, creating the pressing need to establish bioinformatic community standards. Recently, several new search algorithms were released, many of which vary in terms of search strategy, localization system, score cutoffs, and glycan databases, thus warranting a comprehensive comparison of these new programs along with existing programs. Here, we analyzed three common samples: an enriched cell lysate, a mixture of 6 glycoproteins, and a mucin-domain glycoprotein. All raw files were searched with comparable parameters among software and the results were extensively manually validated to compare accuracy and completion of the output. Our results highlight the continued need for manual validation of glycopeptide spectral matches, especially for O-glycopeptides. Despite this, O-Pair outperformed all other programs in correct identification of O-glycopeptides and its localization system proved to be useful. On the other hand, Byonic and pGlyco performed best for N-glycoproteomics; the former was best for proteome-wide searches, but the latter identified more N-glycosites in less complex samples. Overall, we summarize the strengths, weaknesses, and potential improvements for these search algorithms.

**TOC Figure:** 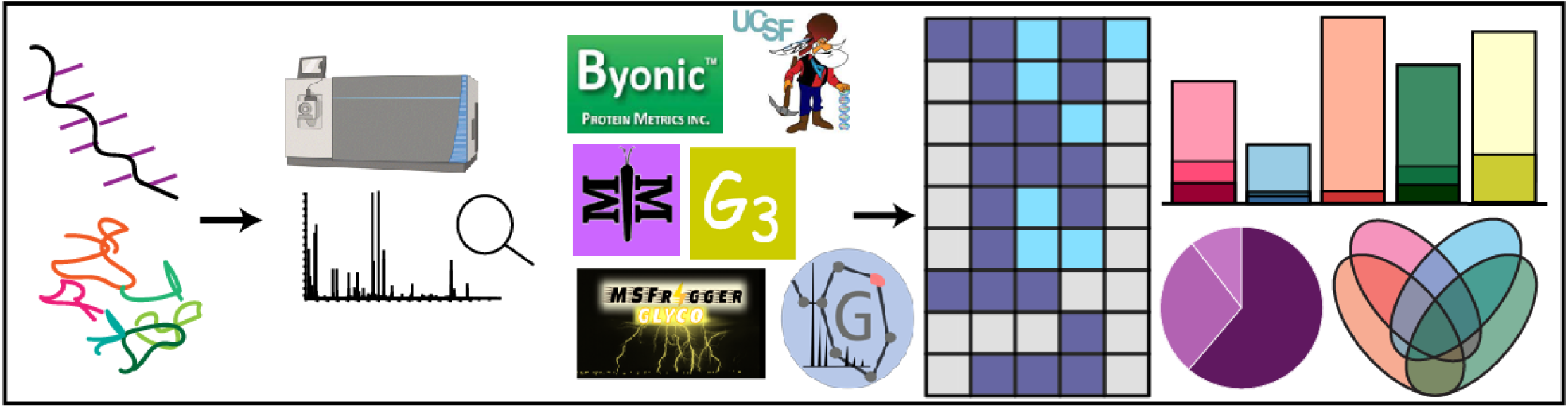

## 1. Introduction

Glycosylation is a post-translational modification (PTM) implicated in viral infection,^1,2^ T cell activation and differentiation,^3^ reproductive health,^4^ and cardiovascular disease.^5^ Changes in glycosylation macrohetereogeneity (site occupancy) and microheterogeneity (distinct glycoforms on one site) are known to be relevant in receptor pattern recognition by influenza A viruses,^6^ interaction of plasma proteins,^7^ cancer,^8,9^ and protein/glycan interactions,^10^ among many others. This highlights the need to accurately detect changes in glycan localization and composition. Mass spectrometry (MS) has emerged as a powerful tool for uncovering the glycomic and glycoproteomic landscape of proteins. While there are five major types of glycosylation, N- and O-glycosylation are the most common. In N-glycosylation, glycans are attached to Asn when the N-X-S/T consensus motif is present.^11^ O-glycans can be attached to any Ser or Thr, regardless of peptide sequence. Mucin domains are regions of dense O-glycosylation with a high content of Pro, Thr, and Ser; within these domains, it is commonly observed that most Ser and Thr residues are modified.^11^

Various computational approaches can be taken to analyze glycoproteomic data. Many of the commonly used search algorithms, such as SEQUEST,^12^ MASCOT,^13^ and Andromeda,^14^ are able to search glycosylation as a variable modification. However, glycopeptide analysis by MS presents challenges that more commonly searched modifications (e.g., deamidation, oxidation) do not have. For instance, collision-based fragmentation produces peptide fragments without the glycan mass, so programs must anticipate the glycan loss to correctly calculate the peptide fragment masses. The most abundant ions in collision-induced glycopeptide MS2s are fragments corresponding to partial glycan losses and/or protonated glycan oxonium ions. Less targeted software can falsely assign these glycan losses as peptide fragments. Additionally, many peaks in the spectrum will remain unassigned, resulting in poor scoring and probability calculations. While it is possible to use most proteomic search software for glycosylation by including it as a variable modification, algorithms that do not consider these challenges will struggle to identify and score glycosylated peptides appropriately. Thus, the field is largely shifting away from the traditional proteomics approach.

To account for these drawbacks, several bioinformatic strategies have been developed with glycosylation in mind. For instance, in the “glycan-first” search method, oxonium ions are used to filter possible glycan structures and, hence, glycan masses.^15^ Then, fragment ions without the glycan mass are searched. Conversely, in the “peptide-first” strategy, fragment ions without the glycan mass shift are searched and fragments are identified based on an “open” or “off-set” search. In an open search, peptide fragment ions are matched in the MS2 regardless of whether the proposed peptide mass matches the mass of the precursor ion in the MS1. In an off-set search, the mass difference between expected fragments and observed fragments is considered as the corresponding glycan mass.^16^

Collisional fragmentation causes difficulties regarding site localization, which is predominantly a concern when investigating O-glycosylation. Given the heterogeneity of glycosylation, if multiple potential sites of O-glycosylation exist, site localization of singly modified peptides is difficult. When the number of plausible modifications in a glycopeptide increases, the possible combinations of glycoforms increases exponentially, introducing an additional level of complexity in software analysis.^17–19^ Normally, collision-based fragmentation only provides total glycan composition and the naked peptide sequence, thus information about multiple glycosites is difficult to ascertain. Some of these challenges can be addressed using electron dissociation (ExD) methods, especially if the peptide sequence is already known; the most common of these is electron transfer dissociation with (EThcD) or without (ETD) supplemental activation. ExD methods can allow for site localization, even if multiple sites of glycosylation exist. That said, multiply O-glycosylated peptides frequently have low charge density. This comes as a consequence of their peptide backbone sequence, which has a high density of uncharged residues and/or sialic acid content, requiring an additional proton for the peptide to be positively charged. Given these issues, glycoproteomic workflows, especially for O-glycoproteomics, often consist of HCD-product dependent-EThcD (HCD-pd-EThcD), where higher energy collisional dissociation (HCD) scans are taken for all precursor ions. Detection of the “HexNAc fingerprint” (i.e., oxonium ions generated from GalNAc or GlcNAc fragmentation) triggers EThcD scans.^18^ How software considers these paired scans (e.g. pairing, merging, or searching them separately) thus affects how glycans are localized.

*MSFragger-Glyco* uses open and mass offset search algorithms to search glycoproteomics data in a single pass, making a fragment ion indexing approach computationally feasible. A single-pass algorithm will read input in one iteration.^20^ In a fragment indexing approach, an array transformation allows constant time querying of fragment indexed ions to run the workload as fast as possible.^21,22^ Only peptides with a possible glycosite and spectra with a user-defined number of oxonium ions are examined for glycan mass offsets.^23^ Importantly, MSFragger-Glyco does not have the capability to localize more than one glycosylation site, which is insufficient in the case of highly modified O-glycoproteins, like those containing mucin domains. MSFragger is hosted in the FragPipe suite which lacks an spectral viewer, but output is supported by Proteomics Data Viewer (PDV).^24^

*O-Pair Search*, part of the MetaMorpheus suite, is the first search algorithm designed for O-glycoproteomic data. The program first performs an open search of HCD spectra, and then uses ions found in EThcD spectra to determine site-specific O-glycan localization(s). Importantly, O-Pair includes glycan localization levels. In Level 1, all possible modifications can be unambiguously assigned with a localization probability of over 0.75. Peptides are assigned as Level 1b if a glycosite has a probability of less than 0.75 or if there is not enough spectral evidence to assign the site. If at least one possible site is unambiguously localized, but others are not, this is denoted as Level 2. Finally, Level 3 identifications represent a confident match of glycopeptide and total glycan mass where no glycosites were able to be assigned unambiguously.^25^

*pGlyco* employs a glycan-first search strategy and a glycan ion-indexing technique to accelerate glycopeptide search speeds. It is capable of handling modified saccharide units,^26,27^ estimating glycan/peptide FDRs, and localizing glycopeptides using stepped collision energy (sce)HCD and ExD spectra. pGlyco merges HCD and ExD spectra into a single spectrum before searching and has several available N- and O-glycan databases for glycopeptide identification. Alternatively, a user can incorporate new glycan databases from GlycoWorkbench.^15^

*GlycReSoft* uses the traditional proteomics approach, where glycans are built in as a variable modification. The algorithm includes a program for building glycan databases from a text file enumerating all compositions and combinatorial constraints describing the space of glycan compositions. It also considers both user and computationally defined glycan modifications and glycopeptide databases. Importantly, it is the only pipeline evaluated here that has the capability to search both glycomic and glycoproteomic data. Additionally, the program averages spectra from the same glycopeptide with different charge states and elution times.^28^

*Protein Prospector* is a search tool whose modification analysis was initially developed for phosphoproteomics and introduced the scoring system later adapted by O-Pair. The program considers modifications, neutral losses, and allows for the comparison of different modification site assignments. It has a viewer incorporated in its pipeline that also displays predicted fragment ions. In previous studies, this algorithm was found to outperform GlycoPAT, GPQust, GlycXTool, among others, in terms of scoring O-glycoproteins and identifying high confidence O-glycopeptides.^29^

Finally, *Byonic* is a commercially available software package for proteomic analysis and allows the user to define an unlimited number of variable modification types. It is currently considered the gold standard in glycoproteomics, though, recent work demonstrated that reported output from Byonic varied widely between groups using the same raw data. Overall, Protein Prospector and Byonic were determined to be high performing for O-glycoproteomics.^29^

In this study, we performed a systematic comparison of six existing and recently released search algorithms for glycoproteomic data. To do so, we generated three samples that represent common glycoprotein preparations with increasing complexity, then searched the resulting MS data files. While most settings were kept consistent between programs, particular settings were tailored to each program given the differences of each algorithm. Finally, we manually validated glycopeptides identified as fully localized by each program and compared the accuracy and completion of output (**Figure S1**). Our results suggest that there is not a “one-size fits all” glycoproteomic software. To be sure, we found that O-Pair outperformed others in O-glycoproteomic analyses, while Byonic performed best in most N-glycoproteomic analyses. Ultimately, we evaluated the accuracy, localization ability, and overall user experience for these bioinformatic platforms.

## 2. Materials and Methods

### Sample preparation

Two digests of recombinant CD43 were performed: Sample A consisted of CD43 digested with trypsin, PNGaseF, and BT4244; Sample B was digested with BT4244, PNGaseF, sialidase, GluC, and trypsin.^30^ For the protein mixture, equal parts (w/w) of fetuin, von Willebrand factor, coagulation factor 12, serotransferrin, CD14, and apolipoprotein E were mixed and digested with trypsin. Finally, lysed JEG3 cells were digested with trypsin and StcE. Further details on digestion conditions and sample preparation can be found in **Supporting Information**.

### Separation and MS

MS analyses were performed using a Thermo Scientific Orbitrap Eclipse Tribrid mass spectrometer coupled to a Dionex Ultimate 3000 UPLC. MS fragmentation settings were set to HCD-pd-EThcD, where EThcD scans were triggered on precursors whose HCD spectra contained 3 out of 9 oxonium fingerprint ions at greater than 5% relative intensity. Detailed separation and acquisition methods can be found in **Supporting Information**.

### Data Analysis

For all searches, database files were downloaded from UniProt. The mucin domain glycoprotein was searched using a database containing the sequence for CD43 (P16150), the protein mixture using a database containing the respective sequences (P12763, P04275, P00748, P02787, P08571, P02649), and the lysate digest was searched with the human proteome (SwissProt, downloaded June 9, 2021). Unless otherwise mentioned, settings for each program were kept at their default values. Fragmentation was set to search HCD and EThcD, when possible. MS1 tolerances were set to 10 ppm and MS2 tolerances were set to 20 ppm. If a multi fragmentation option was not available, EThcD was selected. Carbamidomethyl was set as a fixed modification on Cys, and variable modifications of deamidation (Asn) and oxidation (Met) were allowed. Only identifications under 1% false discovery rate (FDR) were considered using the default decoy generation for each program. For programs unable to process Thermo RAW files, MSConvert (ProteoWizard^31^) was used to generate mzML format. High confidence glycopeptide identifications and localizations were determined based on the developer recommendations.

#### MSFragger

Searches were performed with MSFragger v3.2 using the FragPipe v15.0 pipeline. The respective configurations for hybrid fragmentation methods were used for N- and O-glycopeptide searches. Load setting was set to *Strictrypsin*, which allowed cleavage after Lys/Arg and 5 missed cleavages. The default N- and O-glycan databases contained 181 and 110 structures, respectively (*see* **Supplemental Dataset S1** for a complete list of all glycan databases used). The program was not used for searching the mucin-domain glycoprotein since it expects only one modification per peptide. Modification mode was set to *labile*, and mass offsets were restricted to Asn for N-glycan searches and Ser or Thr for O-glycan searches. PTM-Sheppard was enabled with default configurations from *Glyco Search*. The maximum fragment charge was set to 4 and *Glyco Mode* was enabled. When searching the human proteome, database partitions were set to 50. Identifications with a Hyperscore greater than 10 were considered high confidence and localizations were considered confident when marked as localized by MSFragger.

#### O-Pair

Searches were performed using the MetaMorpheus suite 0.0.319. For O-glycopeptide searches, *O-glycopeptide search* was selected using the default *O-glycan* database, which contained 12 structures. The *N-glycopeptide search* was used for N-glycan searches and the default *N-glycan* database with 182 structures. The protease was set to *nonspecific* with 24 missed cleavages to search the mucin-domain glycoprotein. For the other files, the protease was set to trypsin and 5 missed cleavages were allowed. When searching the human proteome, 50 database partitions were used. Identifications with a Q Value less than 0.01 and localizations assigned as level 1 or 1b were considered high confidence.

#### pGlyco

Searches were performed using pGlyco3.0 rc4. A nonspecific search was used to search CD43 by allowing cleavage C terminally to any amino acid with a maximum of 24 missed cleavages. *The Multi site O-glycan database* containing 3217 glycan configurations was used. Trypsin was selected as the protease for the other files with 5 allowed missed cleavages. The N-human database containing 2922 glycans and the O-human database containing 552 glycans were used to perform separate N- and O-glycan searches. All peptide identifications and localizations with a localization score greater than 5 were considered high confidence.

#### Byonic

Searches were performed using Byonic v4.0.12. For CD43, the cleavage specificity was set to *nonspecific* with zero cleavage sites defined, and any number of missed cleavages were allowed by setting the maximum to -1. The fragmentation type was set to read directly from the scan headers. A maximum of 3 of each glycan was allowed using a database containing the 6 most common O-glycans. Other variable modifications were set as rare, with a maximum of 2 allowed per peptide. For the other files, N- and O-glycans were searched simultaneously, wherein the 9 most common O-glycans were set as common modifications with a maximum of 2 per peptide and 132 N-glycans were set as rare modifications with a maximum of 1 per peptide. For these searches, other variable modifications were set as common, with a maximum of 2 common modifications and 1 rare modification per peptide. For O-glycoproteomic analyses, only EThcD identifications with Delta Mod score greater than 25 were considered for localization. For N-glycoproteomic analyses, initial glycopeptide spectral matches (GSM) filtering for confident peptide sequence (Log Prob > 3 and Score > 300) was applied.

#### GlycReSoft

Files converted to mzML were searched using GlycReSoft v3.0. Using the preset configurations, the *LC-MS/MS Glycoproteomics* option was selected. The maximum charge state was set to 6 and *MSn parameters/averagine* was set to Glycopeptide. A glycan search space was generated using the *Biosynthesis Human N-Glycans* database. Digestion was set to trypsin and a maximum of 2 missed cleavages were allowed. Settings differ from other programs due to difficulties with generation of larger search spaces related to increasing missed cleavages. Due to these difficulties and no option for nonspecific searches, GlycReSoft was not evaluated for O-glycopeptide analysis. Identifications with a q value < 0.01 were considered high confidence.

#### Protein Prospector

Searches were performed in the online suite of Protein Prospector (v 6.3.1) using Batch-Tag Web and mzML files. The instrument was set to *ESI-EThcD-high-res*, precursor charges allowed from 2 to 4, masses set to monoisotopic, and glycopeptide peak filtering was enabled. ExD sialylated variable modifications were allowed. For O-glycopeptide searches, these modifications were set to common and no motif was given. For N-glycopeptide searches, the motif was set to *N[^P][ST]*. The enzyme was set to trypsin with up to 5 missed cleavages, 0 non-specific termini for the mixture and lysate samples, and 2 nonspecific termini for CD43 searches. Identifications with a Pep Score > 7.25 and localizations with SLIP > 6 were considered high confidence.

### Manual Validation

Reported glycopeptides were filtered based on developers’ recommendations. For reported GSMs, the filter parameters were as follows: O-Pair (Q Val < 0.01), Byonic (EThcD spectra only for O-glycopeptides, and log prob > 3 and score > 300 for N-glycopeptides), Protein Prospector (Pep Score > 7.25), GlycReSoft (q < 0.01), and MSFragger (Hyperscore > 10). Then, to generate unique GSMs, duplicates were removed based on full glycopeptide sequence (including glycan composition and localization) for fully localized identifications, and on peptide sequence and glycan mass for non-localized identifications. To filter for software-assigned fully localized GSMs, we used the following parameters: Level 1 and 1b (O-Pair), Localization score > 5 (pGlyco), Delta Mod > 25 for EThcD identifications (Byonic), manual filtering and SLIP > 6 (Protein Prospector), and “MSFragger localized” (MSFragger). For O-glycopeptides, manual validation was performed on all software-assigned fully localized identifications. Here, we generated the expected c/z fragment ions of each glycopeptide and investigated their presence in the corresponding EThcD spectrum. Spectra were averaged if the precursor of the MS2 eluted in the same chromatographic peak. Glycopeptide identifications that only had one possible O-glycosylation site were assigned as localized if manual validation of glycan composition and peptide sequence in the HCD spectrum agreed with what was reported. If the presence of O-glycosylation could not be confirmed for the assigned site using the c/z fragment ions, but its absence on other possible sites was confirmed, the site was assigned as correct. N-glycopeptide sequences and glycan compositions were evaluated using EThcD and HCD spectra. First, we aimed to localize N-glycans in the same manner as O-glycans, by confirming the presence of expected c/z ions. If this was not possible, b/y ions of the assigned peptide sequence were validated in the HCD spectrum. Similarly, glycan composition was validated through glycan losses from the precursor. In some cases, *de novo* sequencing was employed to determine the correct glycopeptide sequence, glycan composition, and localization. Examples of fully localized, incorrectly localized, and glycopeptides unable to be validated for each software can be found in **Figures S2-S9**. Upon initial submission, technical difficulties with MetaMorpheus did not allow for the input of the results file for the N-glycoproteomics search. Similarly, FragPipe lacks a spectral viewer. Therefore, examples from O-Pair and MSFragger are not present in the supplemental figures.

## 3. Results and Discussion

### Glycoproteomic analysis of JEG3 samples

Human epithelial placental choriocarcinoma cell lysate (JEG3) was enriched, digested, and subjected to MS analysis. The N- and O-glycopeptide analyses were performed with six and five search algorithms, respectively (*see* **Methods**). All search algorithm output with annotated validation and total glycopeptide lists from this sample can be found in **Supplemental Dataset S2** and **S3**, respectively.

#### O-glycoproteomics

To account for FDR and correct peptide identification, glycopeptide identifications were filtered to generate unique GSMs. Byonic generated the most unique GSMs (85), followed by MSFragger (80) and Protein Prospector (53) (**Figure 1A**; lightest shade). However, for O-glycosylation, localization of glycans is arguably more important than the number of unique GSMs. Thus, unique GSMs were filtered for localized GSMs per developer recommendations. We found that MSFragger reported the most localized identifications (17), followed by O-Pair (15) and Protein Prospector (13). Interestingly, despite O-Pair reporting nearly half of the unique GSMs when compared to Byonic and MSFragger, a higher percentage of those were considered localized by O-Pair (34%). Remarkably, Byonic only reported 4 localized glycopeptides compared to its initial 85 unique GSMs (**Figure 1A**).

**Figure 1.**
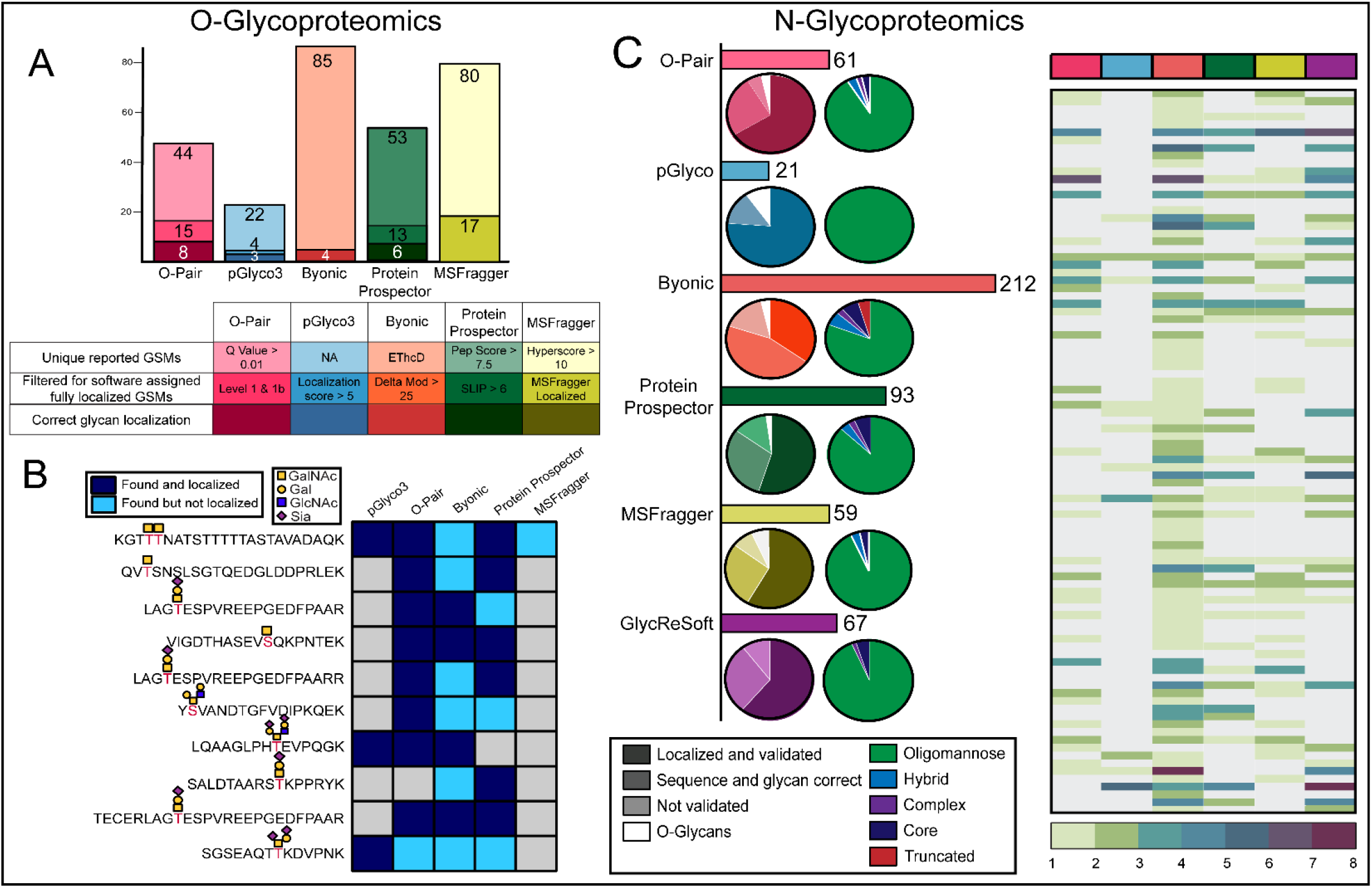
O- and N-glycoproteomic analysis of JEG3. (A) Bar graph representing the different O-glycopeptides identified by five search algorithms. Filtering parameters are described in the table beneath the bar graph: lightest shade - unique glycopeptide identifications; medium shade - software localized identifications; darkest shade - manually validated glycopeptides. (B) True positive O-glycopeptides detected in JEG3. Dark blue signifies that a search algorithm detected and localized associated O-glycan(s) properly, whereas light blue indicates that the peptide sequence was reported but the glycan was not correctly localized by the algorithm. Gray boxes indicate that the glycopeptide was not reported by a search algorithm. (C) Horizontal bar graph represents unique GSMs. The left pie chart demonstrates our manual validation of software localized N-glycopeptides, where the shading follows the same pattern as (A). The white coloration represents O-glycans incorrectly assigned as N-glycans. The pie chart on the right represents N-glycan diversity, where green - oligomannose, blue - hybrid, purple - complex, navy - chitobiose core (“core”), and red - truncated. (D) Glycoform and peptide diversity reported by the six search algorithms. Each row represents an individual peptide sequence and the number of associated glycans reported by each software is indicated by a color (legend at bottom). Gray boxes signify that glycopeptide was not reported by the search algorithm.

Following this, localized GSMs assigned by the algorithms were manually validated. We determined that both O-Pair and Protein Prospector outperformed other programs in terms of their correct assignments and number of glycopeptides reported, where O-Pair correctly localized 53% (8/15), followed by Protein Prospector (46%, 6/13). Despite pGlyco and Byonic having a large percentage correct (75% and 100% respectively), they reported less glycopeptides (3 and 4 localized, respectively). While MSFragger reported 17 localized GSMs, we discovered that 14 of these GSMs were N-glycopeptides assigned as O-glycopeptides and the remaining 3 were incorrectly localized. We then generated a list of true positive O-glycopeptides (**Figure 1B**) and investigated which programs reported the glycopeptides and whether they assigned correct glycan localization. A high agreement between true positives was found in O-Pair, Byonic, and Protein Prospector, where each software reported at least 90% of the glycopeptides. However, O-Pair correctly localized 80% of these peptides (8/10), whereas Byonic and Protein Prospector were more likely to identify the glycopeptide but incorrectly localize the glycosylation (6/10 and 3/10, respectively). Because Byonic does not pair spectra, it requires both identification and localization information to be present in the EThcD spectra, which is not always the case. When applying the recommended filters for confident peptide sequences (Log Prob > 3 and Score > 300), Byonic reported 30 unique GSMs. Unfortunately, none of these were EThcD identifications, highlighting the importance of including information from both ExD and HCD spectra to allow for both identification and correct localization of O-glycopeptides.

#### N-glycoproteomics

Our next goal was to compare how the different software performed in identifying N-glycopeptides, localizing the associated glycans, and detecting different N-glycoforms. As demonstrated by the horizontal bar graph in **Figure 1C**, Byonic reported the most unique GSMs (212), followed by Protein Prospector (93) and GlycReSoft (67). When investigating the accuracy of software-localized GSMs, we found that in more than 75% of cases, we were able to either validate (a) localization with EThcD or (b) peptide sequence and glycan composition using HCD (**Figure 1C**, left pie chart). Despite the lower number of reported GSMs, pGlyco had the highest percentage of correctly localized GSMs (∼75%), but it reported the least number of N-glycopeptides (21). O-Pair had the highest percentage of validated GSMs (∼85%), either via fully localized N-glycans or validated sequence and glycan composition. Additionally, as demonstrated by the white portion of the pie charts in **Figure 1C** (left), Byonic, Protein Prospector, O-Pair, pGlyco, and MSFragger assigned O-glycans as N-glycans in their reported peptide lists. Only GlycReSoft was able to correctly discern between the types of glycans. Programs primarily identified oligomannose N-glycoforms, but Byonic and Protein Prospector detected a larger proportion of diverse glycan types. Protein Prospector detected 3 hybrid, 2 complex, and 5 paucimannose structures which encompassed 12% of total glycan structures. Byonic detected 9 hybrid, 4 complex, 10 paucimannose, and 7 truncated N-glycans, totaling 19% of detected glycoforms (**Figure 1C**, right pie chart). Here, we defined truncated N-glycans as those smaller than the chitobiose core.

Finally, as in the O-glycoproteomic analysis, we generated a list of true positive glycopeptides and investigated which software correctly identified the various glycopeptides and associated glycoforms. Each row in **Figure 1D** represents a different peptide backbone sequence, and the number of different glycans assigned to that sequence is depicted as a heat map. As demonstrated by the high number of colored rows, Byonic detected the most unique glycopeptide sequences (158), with a concomitant high number of isoforms (6 at most). Byonic outperformed other software in N-glycoproteomic searches with Protein Prospector, GlycReSoft, and MSFragger additionally performing well. Here, we found that MSFragger reported a significant number of unique glycopeptides, while Protein Prospector and GlycReSoft found a large number of glycan structures per peptide.

### Glycoproteomic analysis of a complex mixture

After observing significant differences between the identity and accuracy of glycopeptides reported from the JEG3 sample, we prepared a complex mixture of recombinant glycoproteins. Coagulation factor 12 is a glycoprotein essential for blood coagulation^32^ with 2 reported N-glycosites and 27 predicted O-glycosites; serotransferrin is a beta globulin that binds iron for transport^33^ with 3 reported N-glycosites and 2 predicted O-glycosites; von Willebrand factor is an adhesive glycoprotein on the surface of platelets^34^ with 16 reported N-glycosites and 97 predicted O-glycosites; CD14 is a GPI-anchored receptor^35^ with 4 reported N-glycosites and 19 predicted O-glycosites; fetuin is a serum glycoprotein that acts as an inhibitor of soft tissue calcification^36^ with 3 reported N-glycosites and 14 predicted O-glycosites; and apolipoprotein E is a cholesterol carrier that regulates lipid homoeostasis^37^, with 1 reported N-glycosite and 6 predicted O-glycosites. Reported N-glycosites were obtained from Uniprot and O-glycosites were predicted using NetOGlyc4.0.^38^ All search algorithm output with annotated validation and total glycopeptide lists from this sample can be found in **Supplemental Dataset S4** and **S5**, respectively.

Results of this analysis agree with the pattern observed in the JEG sample. In the N-glycoproteomic analyses, none of the algorithms identified all possible N-glycosites annotated in Uniprot. The programs reported a maximum of 2 N-glycosites on coagulation factor 12, 7 on von willebrand factor, 2 on serotransferrin, 1 on fetuin, and 2 on CD14 (**Figure 2**, top). pGlyco and Protein Prospector reported the most N-glycosites (12), followed by Byonic (8). We were able to localize or verify glycan composition and glycan structure in all unique GSMs from Byonic (44), O-Pair (49), and GlycReSoft (22; **Figure S10A**). Similarly, we were able to validate most reported GSMs from MSFragger (16/17) and Protein Prospector (13/15). While pGlyco reported the most unique GSMs (122), we were unable to validate 10 of them.

**Figure 2.**
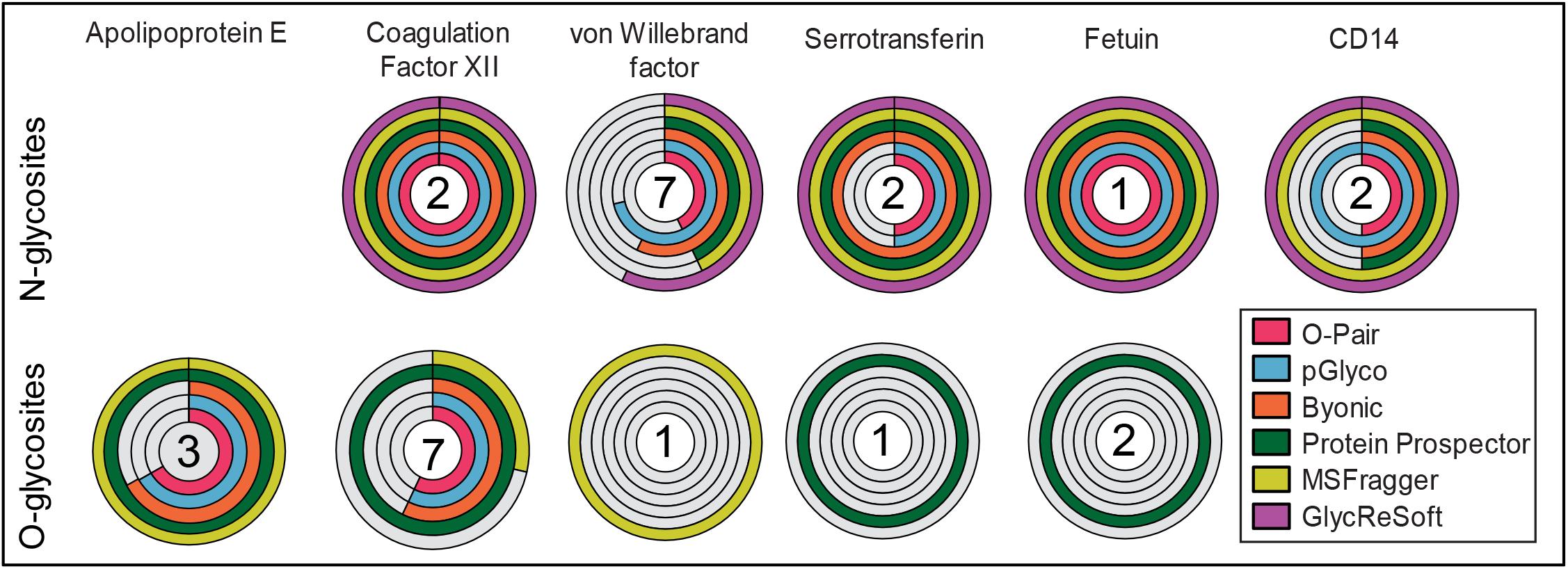
N- and O- glycoproteomic analysis of a glycoprotein mixture. Doughnut plots of the total glycosites found by each software for the 6 proteins present in the glycoprotein mixture. Sites are divided into N- (top) and O- (bottom). The total number of sites identified among all six programs is displayed inside the doughnut plot. As in Figure 1 and depicted in the legend, green represents Protein Prospector, yellow MSFragger, purple GlycReSoft, blue pGlyco, orange Byonic, and pink O-Pair. The maximum number of identified N- and O-sites, respectively, in each of the proteins are as follows: 3 (O-) for apolipoprotein E, 2 and 7 for coagulation factor 12, 7 and 1 for von willebrand factor, 2 and 1 for serotransferrin, 1 and 2 for fetuin, and 2 (N-) for CD14.

In the O-glycoproteomic analyses, only 3 O-glycosites were identified in apolipoprotein E, 7 in coagulation factor 12, 1 in von Willebrand factor and serotransferrin, and 2 in fetuin. Protein Prospector was the only software to identify O-glycosites on fetuin and serotransferrin, and MSFragger exclusively reported O-glycosites of von Willebrand factor (**Figure 2**, bottom). Protein Prospector identified the most O-glycosites (9) followed by O-Pair (7). When performing the O-glycoproteomic analysis, the correctly localized GSMs were as follows: 6/7 for Byonic, 8/15 for O-Pair, 8/13 for pGlyco, 3/19 for MSFragger, and 18/89 for Protein Prospector (**Figure S10B**).

None of the software identified and/or localized all glycosites validated in the protein mixture. However, Protein Prospector identified the most sites (22), followed by pGlyco (18). A combination of the glycopeptides reported by these programs accounted for all of the identified glycosites, which indicates that one might benefit from using more than one software when studying protein mixtures that are less complex than cell lysates. While pGlyco identified very few peptide sequences in the cell lysate, its identifications improved dramatically with the glycoprotein mixture. This may be due to changes in peptide and/or localization scoring with lower protein database complexity. Expected N- and O-glycosites along with sites validated from all algorithms can be found in **Tables S1-S6**.

### O-glycoproteomic analysis of a mucin-domain glycoprotein

CD43 is an extracellular T cell glycoprotein and contains a mucin domain consisting of 235 amino acids. It participates in cell-cell adhesion, signaling, migration, and apoptosis.^39,40^ In our O-glycoproteomic analyses, a total of 51 O-glycosites were verified in the mucin domain of CD43, which is comparable to the 67 sites found previously using a combination of multiple mucinases.^30^ Only core 1 structures were identified, which agrees with previous glycomic analysis.^41^ BT4244 is a mucinase that cleaves N-terminally to glycosylated Ser/Thr, with preference for desialylated core 1 structures.^30^ As expected, due to treatment with BT4244 and sialidase prior to analysis, most glycosylation sites were modified by desialylated core 1 structures. All search algorithm output with annotated validation and total glycopeptide lists from this sample can be found in **Supplemental Dataset S6** and **S7**, respectively.

Normally, using non-specific digestion parameters and allowing for multiple O-glycosites increases search time. However, we observed insignificant differences in search time when using pGlyco, O-Pair, and Protein Prospector; all searches were completed under 5 minutes. In agreement with previous reports,^25^ search time when using Byonic was significantly higher, even when using glycan databases containing only 6 or 9 O-glycan structures.

As in previous sections, identified glycopeptides were filtered and duplicates were removed. O-Pair reported the most unique glycopeptide identifications (460), followed by pGlyco (305) and Byonic (246; **Figure 3A**, light shade). Additionally, O-Pair reported the largest number of localized unique GSMs (143), followed by Byonic (52) and pGlyco (43). After manually validating correctly localized O-glycopeptides, the percentage of accurately reported glycopeptides for each program was found to be 63.46% for Byonic, 48.84% for pGlyco, 48.25% for O-Pair, and 48.57% for Protein Prospector (**Figure 3A**, dark shade). Overall, after manual validation, O-Pair reported the most correctly localized GSMs (69).

**Figure 3.**
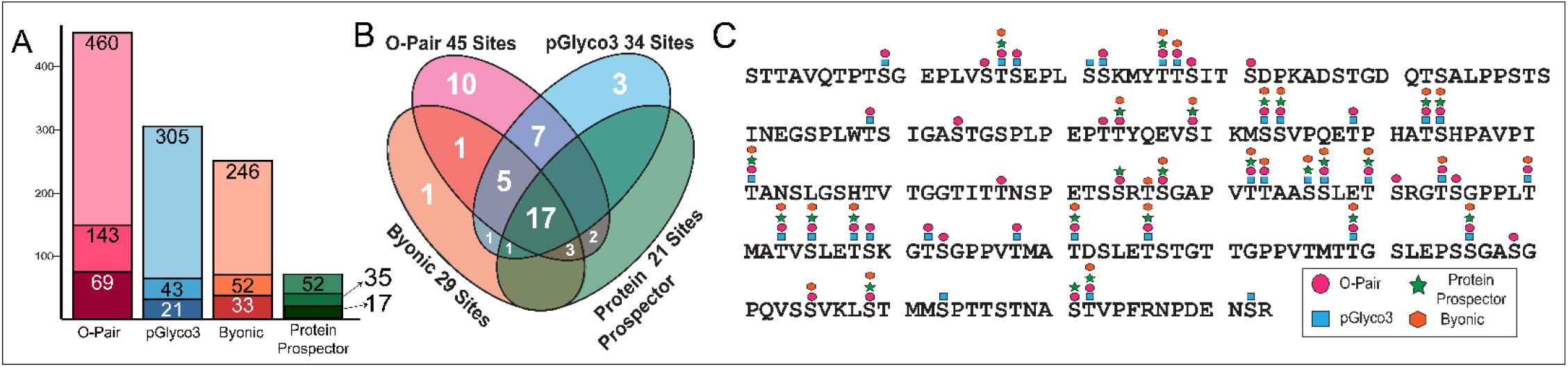
O-glycoproteomic analysis of CD43, a mucin-domain glycoprotein. (A) Bar graph of unique GSMs (light), software-localized GSMs (medium), and manually validated (dark). (B) Venn diagram demonstrating overlap of sites identified between software. Site agreement among the four programs was found to be 17; O-Pair found 10 unique sites, followed by pGlyco (3) and Byonic (1). (C) Peptide sequence of the extracellular mucin-domain of CD43. Identified glycosites are indicated by the presence of a symbol representing the software program that identified the site.

Upon comparing all the total glycosites found by each software, we observed significant differences (**Figure 3B**). In fact, only 33.33% of the total O-glycosylation sites (17/51) were correctly reported by all four search algorithms. Importantly, 14 O-glycosites were reported by only one program, where O-Pair reported 10 unique glycosites, followed by pGlyco (3) and Byonic (1). While most O-glycosylation sites were reported by O-Pair, 6 sites (11.77%) were identified exclusively by other programs. Thus, in targeted analyses of mucin-domain glycoproteins, it may be prudent to use more than one search algorithm to identify the highest number of O-glycosites. More importantly, our results highlight the need for manual validation prior to publishing results in literature, given the low percentage of correctly localized glycopeptide identifications observed among all software. The distribution of reported O-glycosites by the various search algorithms is visualized in **Figure 3C**.

### Summary of strengths for N- and O-glycoproteomic search algorithms

Overall conclusions from our systematic comparison can be found in **Figure 4**. O-Pair performed the best for O-glycoproteomic analysis, having reported the most total glycosites, unique number of glycosites, and the highest percentage of correctly localized glycopeptides. This success can, in part, be attributed to the O-Pair localization scoring system and the 138/144 ratio calculation. Byonic, pGlyco, and Protein Prospector were also evaluated for O-glycoproteomics, each with their own strengths and weaknesses. Both pGlyco and Protein Prospector provide the user with information about potential sites, even if they cannot be fully localized. Additionally, Byonic and Protein Prospector demonstrated a strong ability to identify correct glycan masses and peptide backbones, even though localization was not as strong as in O-Pair. We note that MSFragger and GlycReSoft were incompatible with searching O-glycopeptides, and thus are not recommended for these purposes. Importantly, when applying the developer-recommended filtering parameters for Byonic, most software-assigned localized glycopeptides (i.e., those from EThcD spectra) were filtered out. This is likely because of the low peptide fragment information gathered from these scans. This highlights the need to pair spectra, or to at least consider them, when performing O-glycoproteomic searches. On the other hand, the best performing N-glycoproteomic search algorithm was less clear. While Byonic identified the most unique glycopeptides and the most N-glycan diversity in the JEG3 sample, pGlyco and Protein Prospector identified the most N-glycosites in the protein mixture.

**Figure 4.**
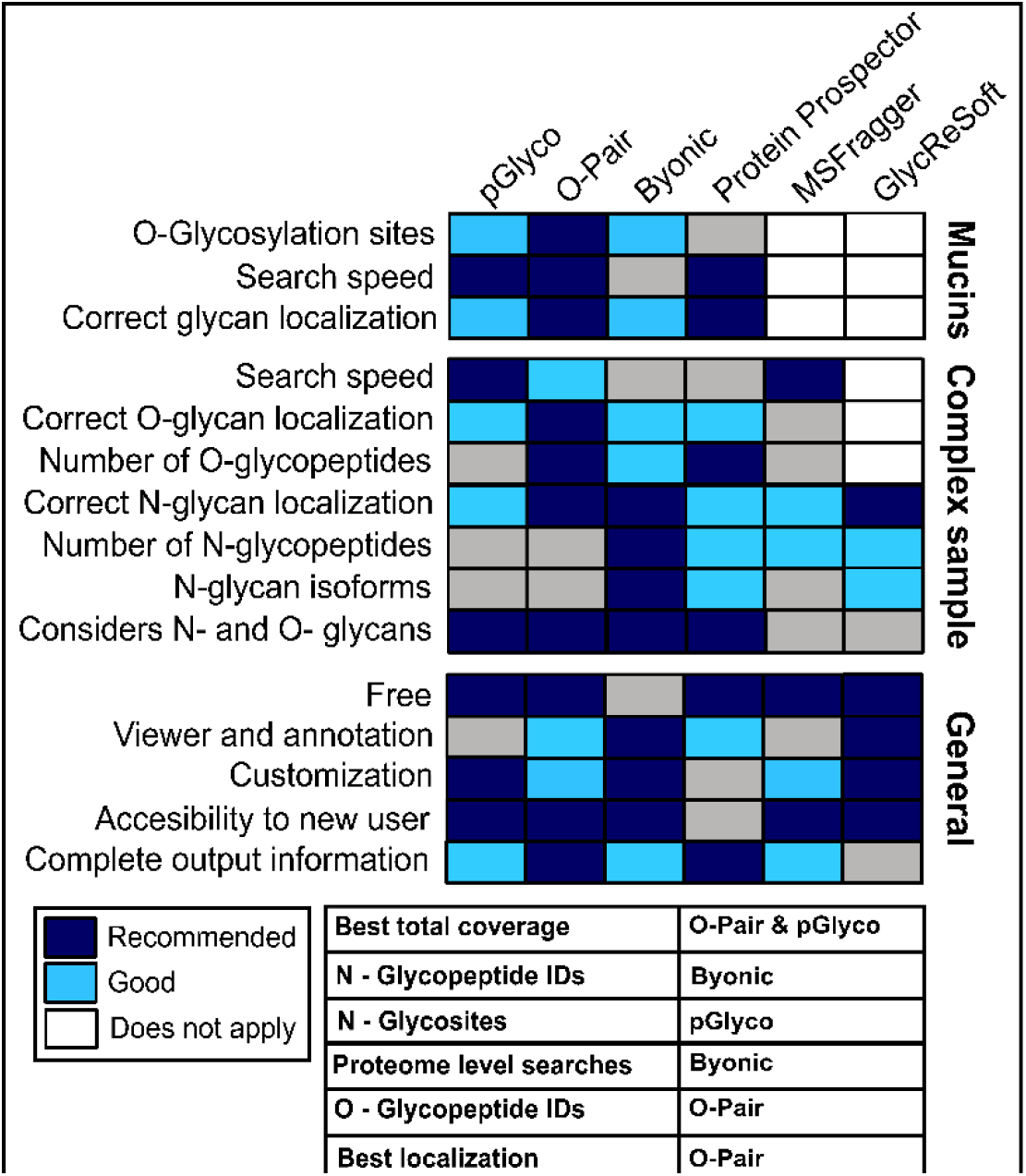
Recommendations for glycoproteomic software. Heat map representing whether the software is recommended (blue) or good (light blue) for a given category. When the category does not apply (e.g., GlycReSoft in O-glycosylation categories) the box is white. If the software outperformed others, it is labeled as recommended; if it yielded good results but did not perform as well as other programs, it is labeled as good.

Several factors must be considered when selecting an ideal search algorithm. For instance, we note that Byonic is the only commercial search algorithm used here, which may be important for some users. Additionally, search speeds are a critical consideration in glycoproteomics, as analyses involving complex PTMs consume significant computing power. As reported previously, most of the programs allowed extremely fast (∼5 min) searches for most samples, whereas Byonic finished after several hours despite using much smaller glycan databases. That said, it is currently possible to perform N- and O-searches simultaneously in Protein Prospector, Byonic, and GlycReSoft, but we were unable to generate a combined N and O-glycopeptide search space in GlycReSoft due to extensive processing time. Therefore, while this option is available among these three, Byonic and Protein Prospector were the only programs to perform successful simultaneous N- and O-glycopeptide searches.

The customizability and user interface are also important considerations when choosing a search algorithm. In this manner, pGlyco, Byonic, MSFragger, and GlycReSoft offered the highest amount of customizability – in terms of the ability to select glycan databases, to add modified glycans, and to change the N- and C-termini cleavage specificity. MSFragger also afforded the ability to customize which HexNAc fingerprint ions, glycan fragmentation, and number of diagnostic ions to consider. While O-Pair offered different termini cleavage options, using these alternatives came at the consequence of lower search performance. For the user interface and viewer/annotation options, Byonic and GlycReSoft were particularly useful for easier data analysis and manual validation. Protein Prospector also had an annotated viewer, but its user interface was the most difficult to use. While it outperformed some other programs in localizations and identifications, the searches had the least amount of customization and could only search one fragmentation type at a time.

Finally, the output between different software varied significantly. As decreed by Minimum Information Required for a Glycomics Experiment (MIRAGE)^42^ and the Human Glycoproteomics Initiative (HGI)^29^, standardized output is a key direction for the field of glycoproteomics. Based on these analyses, we found that the necessary information for validating and reporting glycopeptides and glycosites included: protein identification, retention time, MS2 scan number, fragmentation type, intact glycopeptide m/z, mass error (ppm), peptide sequence, glycan position within the protein and peptide, glycan composition, acknowledgement of the N-X-S/T motif, and 138/144 ratio. O-Pair reported the most complete information, whereas pGlyco was missing the amino acid positions in a protein sequence and GlycReSoft did not report the intact glycopeptide m/z.

## 4. Conclusions

Several challenges are associated with bioinformatic processing of glycoproteomic data. This is largely attributable to the micro- and macroheterogeneity of glycosylation, need to employ multiple dissociation methods, and lability of the glycosidic linkages. Here, we compared some of the most prominent glycoproteomic search algorithms using commonly prepared samples: a mucin-domain glycoprotein, a glycoprotein mixture, and a cell lysate. When analyzing JEG3, we found high O-glycopeptide agreement between Protein Prospector, Byonic, and O-Pair. However, O-Pair was able to assign correct sites for most of the O-glycans while other programs struggled with localization. In our N-glycoproteomic analysis, we found that Protein Prospector, Byonic, and pGlyco outperformed other software in distinct areas: Byonic reported the most unique glycopeptide sequences and associated N-glycans, while pGlyco and Protein Prospector identified the most glycosites in the glycoprotein mixture. In our analysis of CD43, a mucin-domain glycoprotein, O-Pair was the best performing in O-glycopeptide identifications and localization of O-glycosites.

In closing, we found that a “one-size fits all” search algorithm does not currently exist for glycoproteomics. Importantly, the relatively low percentage of correctly assigned identifications highlights the need for thorough manual validation of GSMs prior to reporting glycosylation sites and glycan structures in the literature.

## Acknowledgements

S.A.M. is currently supported by the Yale Science Development Fund and the Yale SEAS/Science Program to Advance Research Collaboration (SPARC). The authors would also like to thank J. Shabanowitz and D.L. Bai for supplies and technical support, R. McCloud for JEG3 cell culture, and C.R. Bertozzi and D.J. Shon for the BT4244 plasmid. Additionally, the authors would like to thank all software developers for their technical support, and Alexandra Steigmeyer for critical reading of this manuscript.

## Notes

### Competing Interest Statement

S.A.M. was formerly a member of the Bertozzi laboratory, which co-produced reference number 25.

