## Supplemental Information for "A systematic comparison of current bioinformatic tools for glycoproteomics data"

**This PDF file includes:**

Supplementary Materials and Methods

Figures S1 to S10

Tables S1 to S6

**Additional supplementary materials not included in this document:**

Supplemental Dataset S1 - Glycan Databases

Supplemental Dataset S2 – Output and annotated validation for JEG3

Supplemental Dataset S3 – Glycopeptides found in JEG3

Supplemental Dataset S4 – Output and annotated validation for mixture

Supplemental Dataset S5 – Glycopeptides found in mixture

Supplemental Dataset S6 – Output and annotated validation for CD43

Supplemental Dataset S7 – Glycopeptides found in CD43

#### Supplementary Materials and Methods

**Sample preparation:** All human recombinant proteins were acquired from R&D Systems, unless specified. Solvents were purchased from Thermo Fisher and LC-MS grade unless specified. Buffer A (0.1% formic acid (28905) in Pierce water LC-MS grade (51140)), Buffer B (80% acetonitrile (ACN, Honeywell, UN1648) in 0.1% formic acid), and Buffer C (40% ACN in 0.1% formic acid) were used throughout sample preparation.

*CD43*: Sample A consisted of CD43 digested with trypsin, PNGaseF and BT4244, while sample B was digested with BT4244, PNGaseF (New England BioLabs), sialidase (SialEXO, Genovis), GluC (Promega, sequencing grade) and trypsin (Promega, sequencing grade). In sample A, 1 µg of CD43 (P16150 - 9680-CD-050) was reconstituted in 10 µL of LC-MS water and 2 µL of dilute BT4244 (0.09µg/µL in LC-MS water); similarly, sample B consisted of 1 µg of CD43 reconstituted in 9 µL of LC-MS water, 2 µL of dilute BT4244, and 1 µL of dilute SialEXO (2.5% by volume in LC-MS water). Samples were incubated at 37 °C overnight and centrifuged for 2 min at 13000 *x g*. For PNGaseF digestion, 7 µL of 50 mM ammonium bicarbonate (A6141, Sigma), and 1 µL of dilute PNGaseF (2% by volume in LC-MS water) were added, followed by incubation for 8 hr at 37 °C. Protease digestion was then performed by adding 1 µL of LC-MS water and 1 µL of 0.05 µg/µL trypsin to sample A. Sample B was reacted with 1 µL of 0.05 µg/µL GluC and 1 µL of 0.05 µg/µL trypsin. Digestion was incubated overnight at 37 °C and quenched by adding 0.5 µL of formic acid.

*Protein Mixture:* The protein mixture consisted of equal parts of alpha-2-HS-glycoprotein (fetuin) (P12763 - 1725-PI), von Willebrand factor (P04275 - 2764-WF), coagulation factor XII (P00748 - 1473-SE), serotransferrin (P02787 - 2914-HT), monocyte differentiation antigen (CD14) (P08571 - 383-CD/CF) and apolipoprotein E (P02649 - 3520-AR). Here, 2 µg of each protein was combined and diluted in 60 µL of 50mM ammonium bicarbonate. To reduce and alkylate, 3 µL of 25 mM dithiothreitol (Sigma Aldrich, D0632) in LC-MS water was added to the sample, followed by incubation at 56 °C for 20 min. The samples were held at RT for 10 min to avoid overalkylation followed by the addition of 3 µL of 35 mM of iodoacetamide (Sigma Aldrich, I1149) for 15 min in the dark at RT. After diluting with 30 µL of 50 mM ammonium bicarbonate, 5 µL of 0.05 µg/µL trypsin was added, reacted overnight at 37 °C, and the reaction was quenched by adding 1 µL of formic acid.

*JEG3:* JEG-3 cells were acquired from ATCC (HTB-36) and cultured according to standard protocols in Dulbecco's modified Eagles medium with 10% fetal bovine serum. Cells were lysed by rotation at 4 °C for one hr in a buffer containing 10 mM Tris pH 8 (Bio-Rad Labs, 1610719), 150 mM NaCl (Sigma Aldrich, S9888), 1% Triton X-100 (Sigma Aldrich, X100), 100U/mL Benzonase Nuclease (EMD Millipore, 70746), 1% Halt Protease (Thermo Scientific, 78440) and phosphatase inhibitor cocktail (Thermo Scientific, 4906845001) followed by 3x 5 sec pulses with a probe sonicator. Cell debris was pelleted by centrifugation at 20,000 *x g* at 4 °C for 20 min. The supernatant was removed, and protein concentration was determined by a BCA assay (Thermo Fisher, 23225). Lysate (250 µg) was subjected to a chloroform/methanol extraction to remove the detergents and protease inhibitors. After reconstitution in 100 µL of 50 mM ammonium bicarbonate, the sample was digested overnight at 37 °C at a 1:50 ratio with metalloprotease StcE. The sample was then reduced in 1 mM DTT at 60 °C for 30 min and alkylated by reacting with 2 mM IAA at RT in the dark. Trypsin/LysC (1.5 µg) was added and allowed to digest overnight on a shaker at 37 °C. The reaction was quenched using 1 µL formic acid and desalted on Strata X-33 cartridges (Phenomenex). Finally, a HILIC enrichment was performed on 50 µg of material, dried and reconstituted in 0.1% formic acid.

**Sample Cleanup:** *C*ontaminants were removed using Strata-X 10 mg SPE C18 1 mL tubes. Here, columns were washed with ACN and equilibrated with buffer A. Samples were diluted to 300 µL in buffer A, transferred to the columns, rinsed with buffer A, and eluted with 300 µL of buffer B or C. Solvent was evaporated in a vacuum concentrator (Labconco) and samples were resuspended in buffer A to reach a final concentration of 0.2 µg/µL. CD43 desalting was performed with 1 mL wash volumes and eluted using buffer B. For all other samples, columns were washed with 0.5 mL of ACN and equilibrated with 0.5 mL of buffer A, rinsed with 300 µL of buffer A 2x, and eluted using 300 µL of Solvent C.

*Separation and MS:* Measurements were acquired in a Thermo Scientific Orbitrap Eclipse Tribrid Mass Spectrometer coupled to a Dionex Ultimate 3000 UPLC. 300ng of samples were loaded from the autosampler onto a C18 trap column (Acclaim PepMap -100µm by 2cm, 5 µm particle size, 100 Å pore size) using Solvent A, which was then washed with loading pump solvent (2% ACN 0.1% formic acid). The trap column was then switched to an EASY-Spray column packed with C18 PepMap material for gradient elution. The column temperature was held at 40 °C using the EASY-Spray ionization source, and samples were eluted at a constant flow rate of 0.3 µg/µL. For the CD43 samples, the flow gradient consisted of 0 to 95% Buffer B in 33 min with a total duration of 60 min. For the remainder of samples, the method used a flow gradient of 0 to 95% Buffer B in 70 min with a total duration of 90 min.

MS measurements were performed in positive ion mode. MS1 scans were acquired in the Orbitrap at 120,000 resolution full width at half maximum (FWHM) at 400 m/z, an automatic gain control (AGC) target of 3e4, and an m/z scan range of 300-1500 m/z. Dynamic exclusion was set to exclude after 3 times for a repeat duration of 10 and an exclusion duration of 10. Charge states 2-6 were filtered for fragmentation, where MS2s were generated with a 3 s duty cycle. HCD was performed on all selected precursor masses with an isolation window of 2 m/z, 30% collision energy, detection in the Orbitrap with a resolution of 15,000 FWHM at 400 m/z, maximum injection time of 50 ms, and an AGC target of 1e4 ions. An EThcD scan was triggered if the HCD scan contained 3 out of 8 oxonium fingerprint ions (126.055, 138.055, 144.07, 168.065, 186.076, 204.086, 274.092, and 292.103) at greater than 5% relative intensity. EThcD used calibrated charge-dependent ETD times, maximum injection time was set to 200 ms, and supplemental activation was set to 20%. Ions were detected in the Orbitrap with a resolution of 30,000 FWHM at 400 m/z.

**Data Availability:** RAW files and search results have been uploaded to PRIDE repository Data are available via ProteomeXchange with identifier PXD032283.

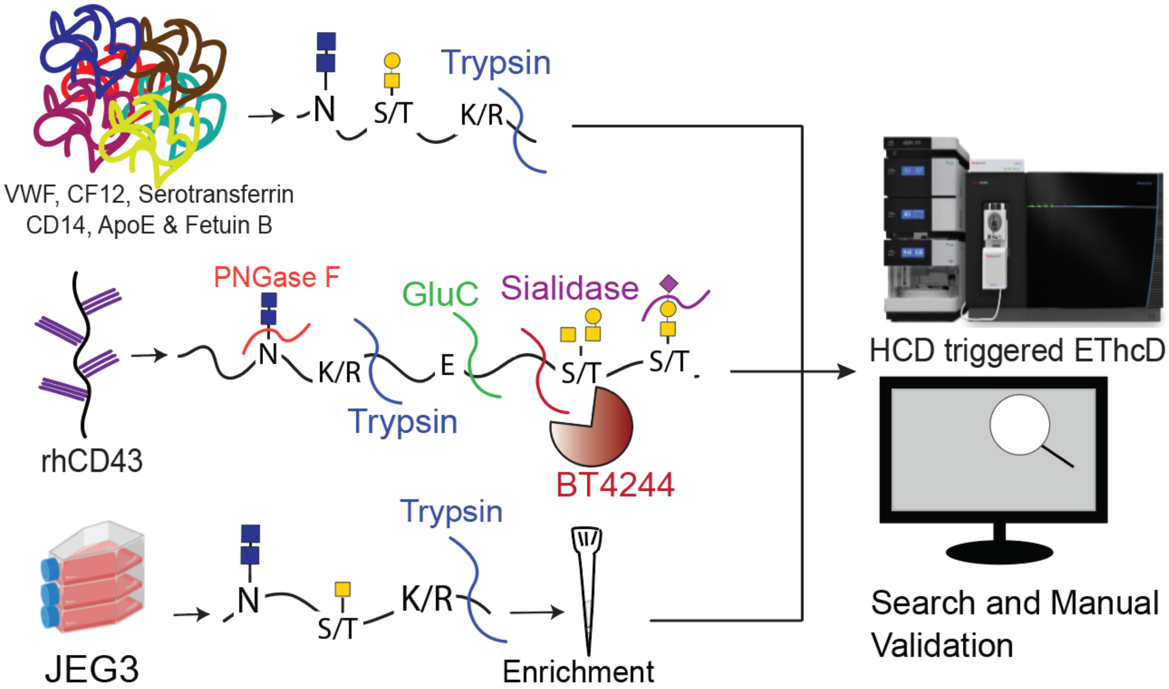


**Figure S1. Workflow of experiments and analyses performed.** Three samples were analyzed, each with various levels of glycosylation. JEG3 cells were lysed, digested with trypsin, and HILIC enriched; a glycoprotein mixture was digested with trypsin; and CD43 was digested with PNGaseF, +/- sialidase, BT4244, trypsin, and +/- GluC. Following digestion, samples were separated and subjected to MS analysis, where an HCD-product dependent-EThcD workflow was applied. Finally, RAW files were analyzed with all search algorithms and software-localized GSMs were manually validated.

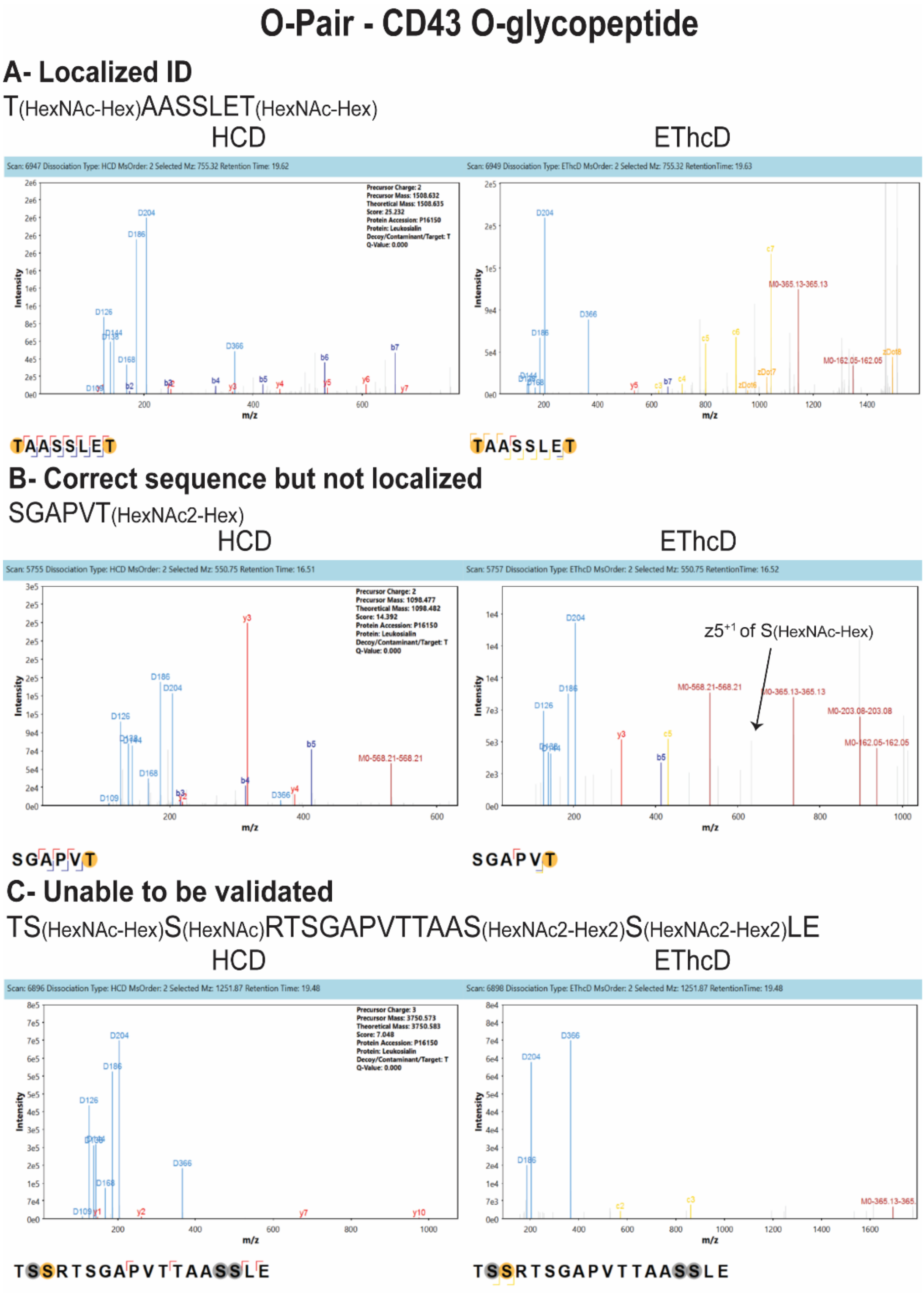


**Figure S2.** Examples of O-glycopeptide localization in O-Pair. (A) shows a completely localized GSM; (B) a GSM with correct sequence but incorrect glycan localization, and (C) a GSM with insufficient localization and sequence information. HCD spectrum are shown on the left and the EThcD are on the right. In (A), HCD fragments supported the correct peptide sequence (TAASSLET) and ETD fragments allowed for complete localization of both glycosites. In (B), peptide sequence (SGAPVT) was confirmed in the HCD. Upon manual validation, we found fragment ion $z_5^{1+}$ corresponding to S(HexNAc-Hex). Therefore, the glycopeptide localization should be S(HexNAc-Hex)GAPVT(HexNAc). Finally, in (C), very little peptide fragmentation corresponding to the proposed peptide was seen in the HCD spectrum. Similarly, there was very little glycopeptide fragmentation in the EThcD spectrum. Here, we could not confidently agree with the proposed peptide. All spectra in this figure were taken from CD43, and annotated spectra were generated using the viewer available in the software suite, MetaDraw.

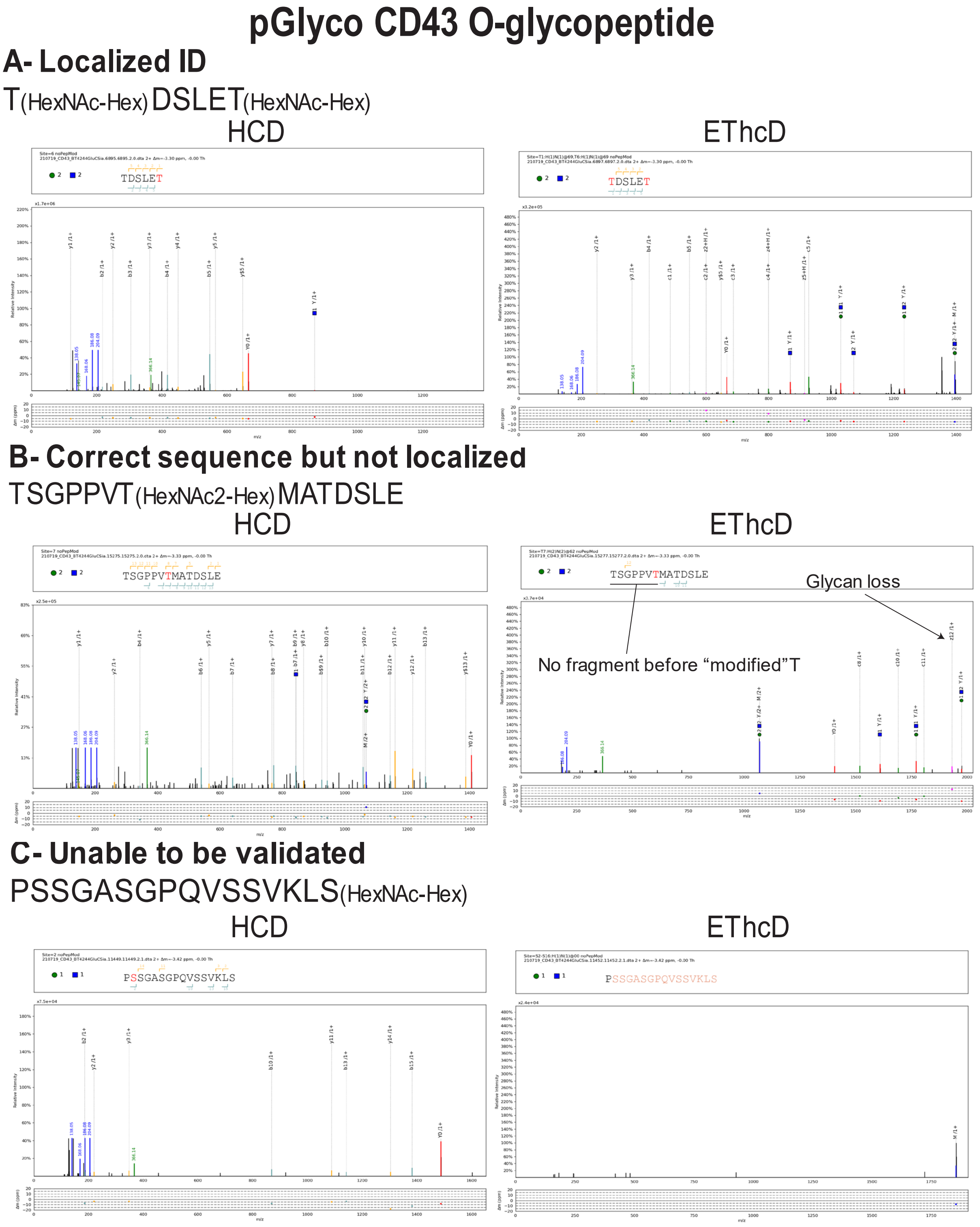


**Figure S3.** Examples of O-glycopeptide localization in pGlyco3. (A) shows a completely localized GSM; (B) a GSM with the correct sequence but incorrect glycan localization, and (C) a GSM with insufficient localization and sequence information. HCD spectrum are shown on the left and the EThcD on the right. (A) HCD fragments supported the correct peptide sequence (TDSLET), and ETD fragments allowed for complete localization of both glycosites. (B) Peptide sequence (TSGPPVTMATDSLE) was confirmed using the HCD spectrum. Upon manual validation, no ETD fragment ions were observed before the assigned glycosite; thus, the Thr could not be validated as modified. In (C), very little peptide fragmentation corresponding to the proposed peptide was seen in the HCD spectrum. Similarly, there was very little glycopeptide fragmentation in the EThcD spectrum. Here, we could not confidently agree with the proposed peptide. All spectra in this figure were taken from CD43 and annotated spectra were generated using gLabel, from pFindStudio.

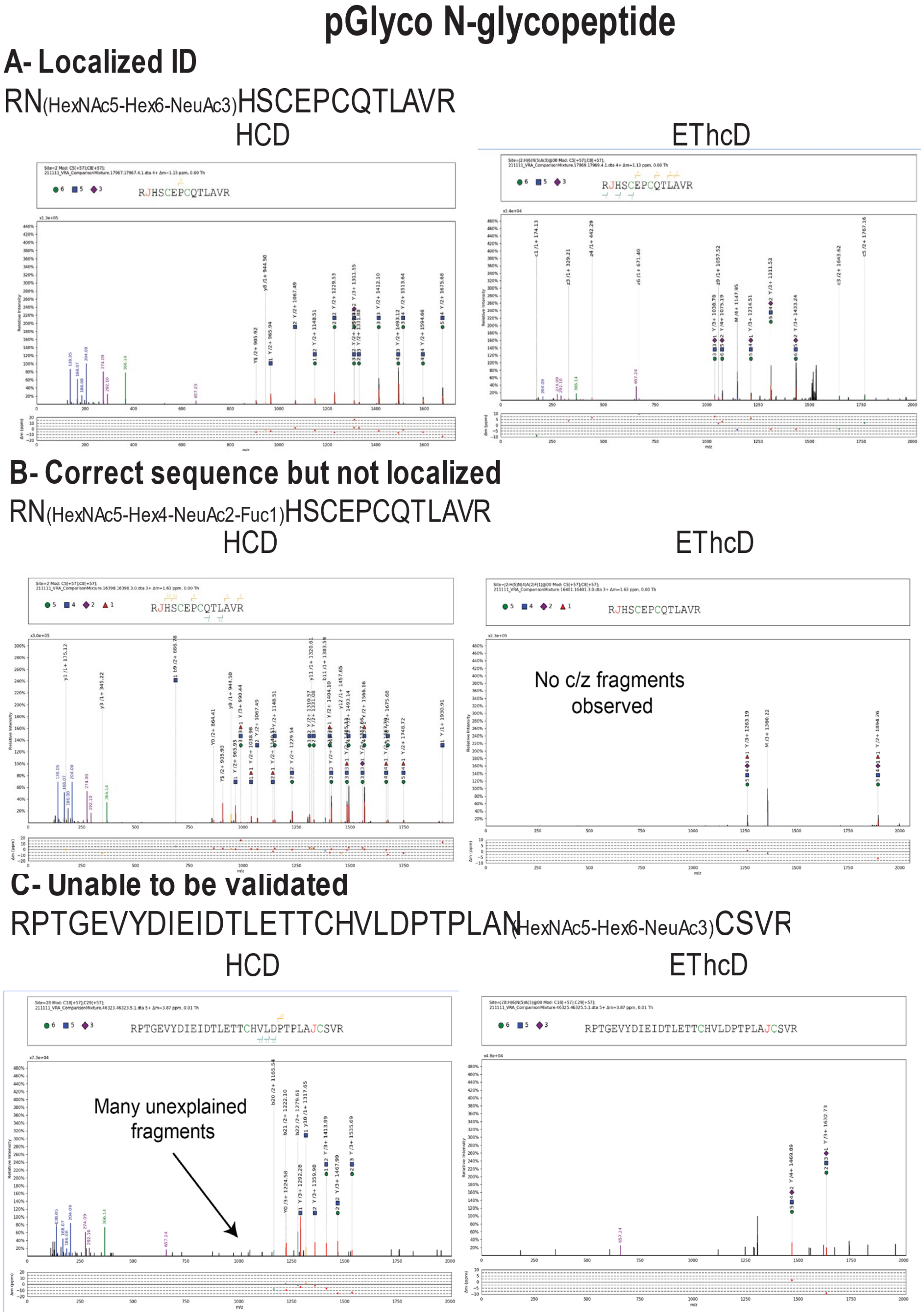


**Figure S4.** Examples of N-glycopeptide localization in pGlyco3. (A) shows a completely localized GSM; (B) GSM with the correct sequence but incorrect glycan localization, and (C) a GSM with insufficient localization and sequence information. HCD spectrum are shown on the left and the EThcD on the right. In (A), HCD fragments supported the correct peptide sequence (RNHSCEPCQTLAVR), and ETD fragments allowed for complete localization of the N-glycosite. In (B), peptide sequence (RNHSCEPCQTLAVR) was confirmed using the HCD. However, no ETD fragment ions were observed; thus, we could not rule out the possibility that the 2 possible O-glycosites were modified. Therefore, there were 2 plausible O-glycosites and 1 N-glycosites. Without ETD information, sites could not be confidently localized. Finally, in (C), many unexplained peptide fragments were observed in the HCD spectrum. Similarly, we observed very little glycopeptide fragmentation in the EThcD spectrum. Thus, we could not confidently agree with the proposed peptide. All examples in this figure were taken from the glycoprotein mixture sample and annotated spectra were generated using gLabel, from pFindStudio.

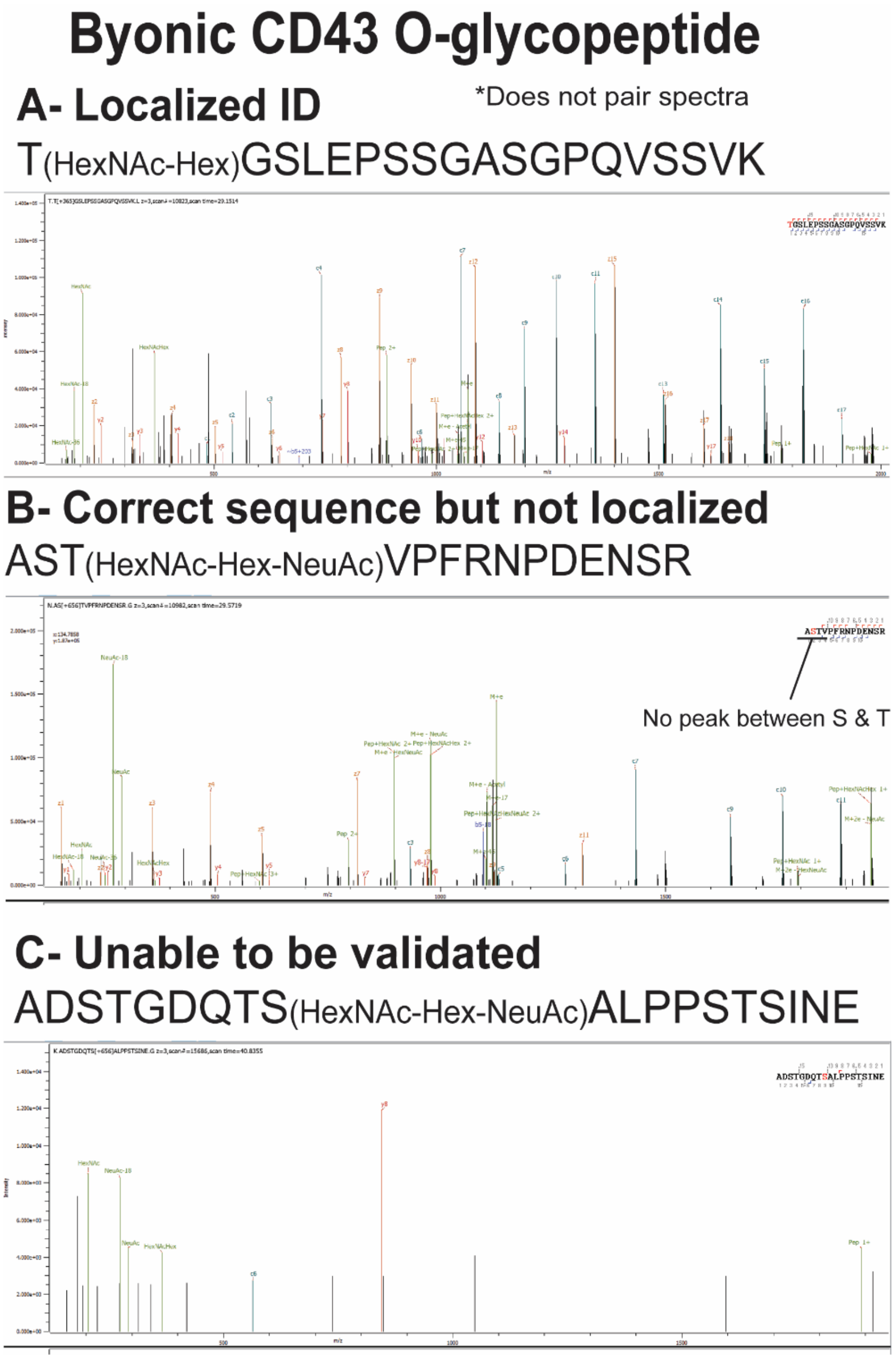


**Figure S5.** Examples of O-glycopeptide localization in Byonic. (A) shows a completely localized GSM; (B) GSM with the correct sequence but incorrect glycan localization, and (C) a GSM with insufficient localization and sequence information. Since Byonic does not pair spectra, only the EThcD spectrum are shown. In (A), EThcD fragment ions confirmed both peptide sequence and glycan localization. (B) shows that peptide sequence was confirmed in the EThcD fragmentation, but no c/z ions were observed between the subsequent Ser and Thr, and therefore, the glycan was not localized to either. Finally, in (C), very little peptide fragmentation corresponding to the proposed peptide was seen in the EThcD spectrum. All spectra in this figure were taken from CD43, and annotated spectra were generated using the PMI Byonic viewer.

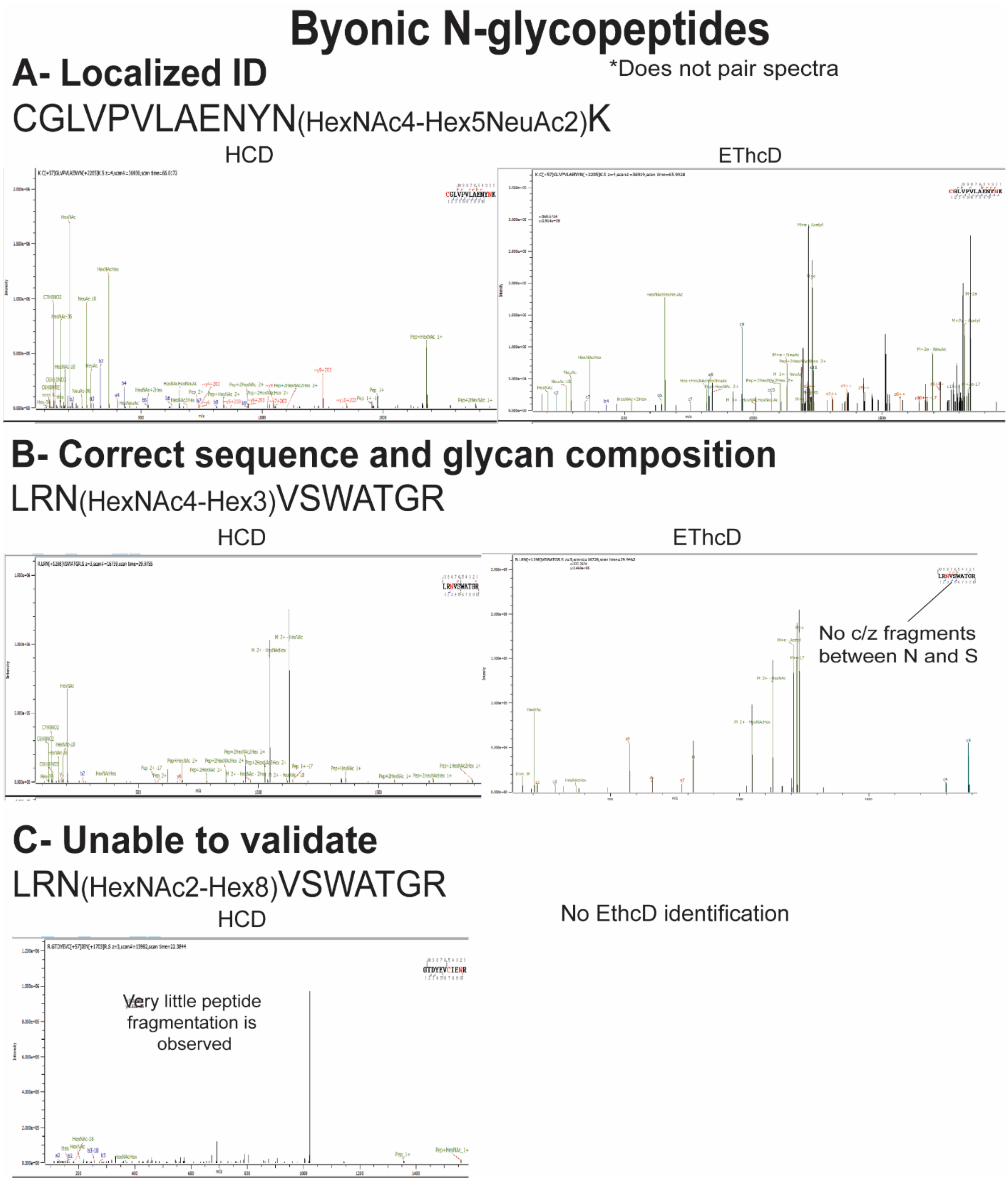


**Figure S6.** Examples of O-glycopeptide localization in Byonic. (A) shows a completely localized GSM; (B) GSM with the correct sequence but incorrect glycan localization, and (C) a GSM with insufficient localization and sequence information. HCD spectrum are shown on the left and the EThcD on the right. In (A), ETD fragment ions confirmed glycan localization and b/y fragment ions confirmed sequence in the HCD scan. In (B), peptide sequence was confirmed in the HCD scan, but fragments were not observed between the N and S, and therefore, the glycan could not be fully localized. Finally, in (C), very little peptide fragmentation corresponding to the proposed peptide was seen in the HCD spectrum, and due to the low abundance of the oxonium ions, an EThcD scan was not triggered. A and B were taken from the mixture sample whereas C was taken from the JEG3 sample. Annotated spectra were generated using the PMI Byonic viewer.

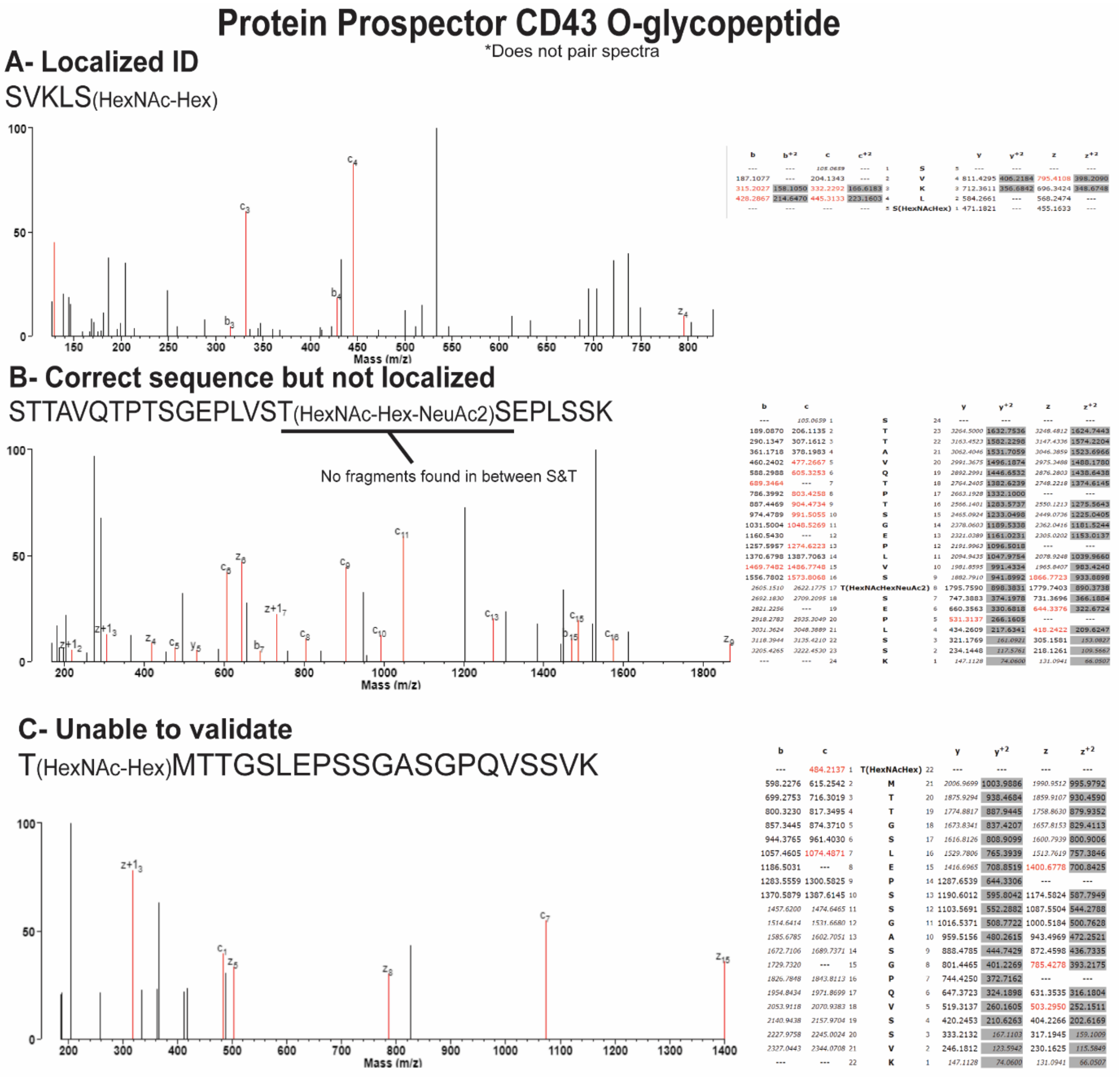


**Figure S7.** Examples of O-glycopeptide localization in Protein Prospector. (A) shows a completely localized GSM; (B) GSM with the correct sequence but incorrect glycan localization, and (C) a GSM with insufficient localization and sequence information. Since Protein Prospector does not consider more than one fragmentation type, only the EThcD spectrum are shown. (A) EThcD fragment ions confirmed both peptide sequence and glycan localization. (B) Here, the peptide sequence was confirmed using ETD fragments, but fragments were not detected between the assigned Thr and the adjacent Ser, thus could not be completely localized. (C) Very little peptide fragmentation corresponding to the proposed peptide was seen in the EThcD spectrum. All examples in this figure correspond to CD43, and annotated spectra were generated using the viewer available in the Protein Prospector software suite.

### Protein Prospector Mixture N-glycopeptides

*Does not pair spectra

#### A- Localized ID

CGLVPVLAENYN(HexNAc4-Hex5-NeuAc2)K

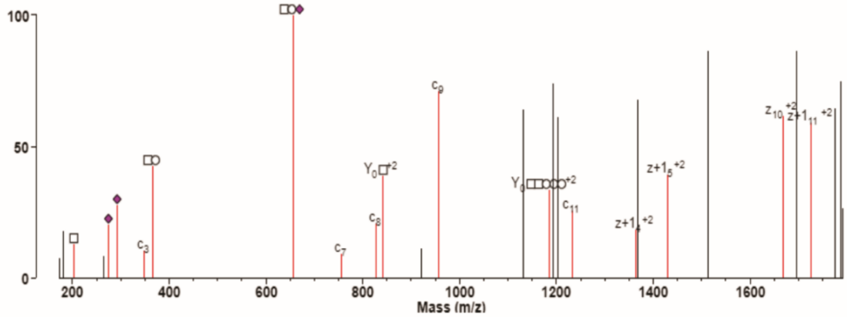


| b | c | # | | # | y | $y^{+2}$ | z | $z^{+2}$ |
|---|---|---|---|---|---|---|---|---|
| --- | 178.0645 | 1 | C(Carbamidomethyl) | 13 | --- | --- | --- | --- |
| 218.0594 | 235.0859 | 2 | G | 12 | 3521.4933 | 1761.2503 | 3505.4746 | 1753.2409 |
| 331.1435 | 348.1700 | 3 | L | 11 | 3464.4719 | 1732.7396 | 3448.4531 | 1724.7302 |
| 430.2119 | --- | 4 | V | 10 | 3351.3878 | 1676.1975 | 3335.3691 | 1668.1882 |
| 527.2646 | 544.2912 | 5 | P | 9 | 3252.3194 | 1626.6633 | --- | --- |
| 626.3330 | 643.3596 | 6 | V | 8 | 3155.2666 | 1578.1369 | 3139.2479 | 1570.1276 |
| 739.4171 | 756.4437 | 7 | L | 7 | 3056.1982 | 1528.6027 | 3040.1795 | 1520.5934 |
| 810.4542 | 827.4808 | 8 | A | 6 | 2943.1141 | 1472.0607 | 2927.0954 | 1464.0513 |
| 939.4968 | 956.5234 | 9 | E | 5 | 2872.0770 | 1436.5421 | 2856.0583 | 1428.5328 |
| 053.5397 | 1070.5663 | 10 | N | 4 | 2743.0344 | 1372.0209 | 2727.0157 | 1364.0115 |
| 216.6031 | 1233.6296 | 11 | Y | 3 | 2628.9915 | 1314.9994 | 2612.9728 | 1306.9900 |
| 3535.4184 | 3552.4450 | 12 | N(HexNAc4Hex5NeuAc2) | 2 | 2465.9282 | 1233.4677 | 2449.9094 | 1225.4584 |
| --- | --- | 13 | K | 1 | 147.1128 | 74.0600 | 131.0941 | 66.0507 |

#### B- Correct sequence but not localized

N(HexNAc4-Hex5-Fuc-NeuAc)VSCPQLEVPVCPSGFQLSC

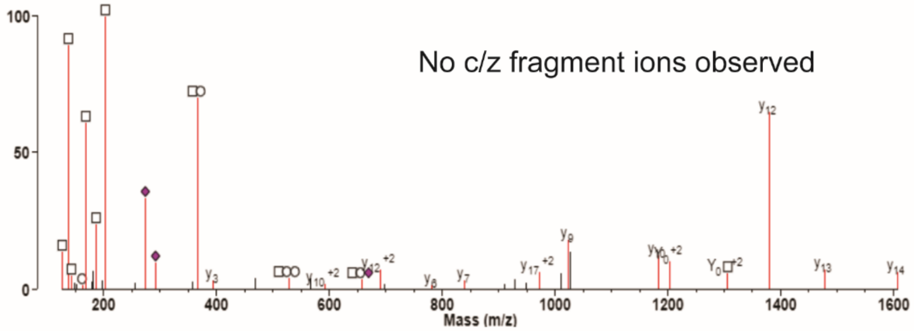


| b | c | # | | # | y | $y^{+2}$ | z | $z^{+2}$ |
|---|---|---|---|---|---|---|---|---|
| --- | 2191.8117 | 1 | N(HexNAc4Hex5FucNeuAc) | 21 | --- | --- | --- | --- |
| 2273.8536 | 2290.8801 | 2 | V | 20 | 2292.0821 | 1146.5447 | 2276.0634 | 1138.5353 |
| 2360.8856 | 2377.9121 | 3 | S | 19 | 2193.0137 | 1097.0105 | 2176.9950 | 1089.0011 |
| 2520.9162 | --- | 4 | C(Carbamidomethyl) | 18 | 2105.9817 | 1053.4945 | 2089.9630 | 1045.4851 |
| 2617.9690 | 2634.9955 | 5 | P | 17 | 1945.9510 | 973.4792 | --- | --- |
| 2746.0276 | 2763.0541 | 6 | Q | 16 | 1848.8983 | 924.9528 | 1832.8795 | 916.9434 |
| 2859.1116 | 2876.1382 | 7 | L | 15 | 1720.8397 | 860.9235 | 1704.8210 | 852.9141 |
| 2988.1542 | 3005.1808 | 8 | E | 14 | 1607.7556 | 804.3815 | 1591.7369 | 796.3721 |
| 3087.2226 | --- | 9 | V | 13 | 1478.7130 | 739.8602 | 1462.6943 | 731.8508 |
| 3184.2754 | 3201.3020 | 10 | P | 12 | 1379.6446 | 690.3259 | --- | --- |
| 3283.3438 | 3300.3704 | 11 | V | 11 | 1282.5919 | 641.7996 | 1266.5731 | 633.7902 |
| 3443.3745 | --- | 12 | C(Carbamidomethyl) | 10 | 1183.5234 | 592.2654 | 1167.5047 | 584.2560 |
| 3540.4272 | 3557.4538 | 13 | P | 9 | 1023.4928 | 512.2500 | --- | --- |
| 3627.4593 | 3644.4858 | 14 | S | 8 | 926.4400 | 463.7237 | 910.4213 | 455.7143 |
| 3684.4807 | 3701.5073 | 15 | G | 7 | 839.4080 | 420.2076 | 823.3893 | 412.1983 |
| 3831.5491 | 3848.5757 | 16 | F | 6 | 782.3865 | 391.6969 | 766.3678 | 383.6875 |
| 3959.6077 | 3976.6343 | 17 | Q | 5 | 635.3181 | 318.1627 | 619.2994 | 310.1533 |
| 4072.6918 | 4089.7183 | 18 | L | 4 | 507.2595 | 254.1334 | 491.2408 | 246.1240 |
| 4159.7238 | 4176.7504 | 19 | S | 3 | 394.1755 | 197.5914 | 378.1568 | 189.5820 |
| 4319.7545 | 4336.7810 | 20 | C(Carbamidomethyl) | 2 | 307.1435 | 154.0754 | 291.1247 | 146.0660 |
| --- | --- | 21 | K | 1 | 147.1128 | 74.0600 | 131.0941 | 66.0507 |

#### C- Unable to validate

CGLVPVLAENYN(HexNAc4-Hex6-Fuc2)KSDNCEDTPEAGYFAIAVVKK

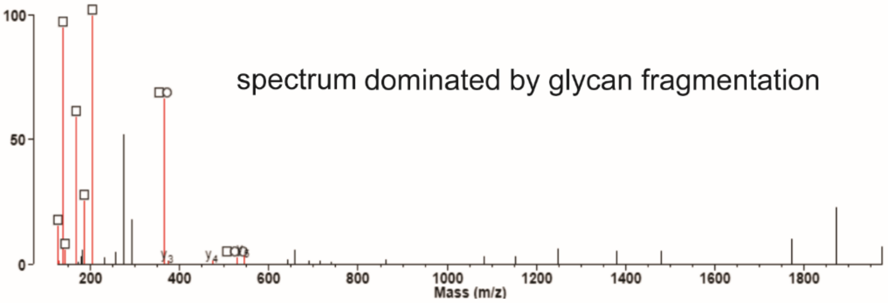


| b | $b^{+2}$ | c | $c^{+2}$ | # | | # | y | $y^{+2}$ | z | $z^{+2}$ |
|---|---|---|---|---|---|---|---|---|---|---|
| --- | --- | 178.0645 | --- | 1 | C(Carbamidomethyl) | 33 | --- | --- | --- | --- |
| 218.0594 | --- | 235.0859 | --- | 2 | G | 32 | 5588.4916 | 2794.7494 | 5572.4729 | 2786.7401 |
| 331.1435 | --- | 348.1700 | --- | 3 | L | 31 | 5531.4701 | 2766.2387 | 5515.4514 | 2758.2293 |
| 430.2119 | --- | --- | --- | 4 | V | 30 | 5418.3861 | 2709.6967 | 5402.3674 | 2701.6873 |
| 527.2646 | --- | 544.2912 | --- | 5 | P | 29 | 5319.3177 | 2660.1625 | --- | --- |
| 626.3330 | --- | 643.3596 | --- | 6 | V | 28 | 5222.2649 | 2611.6361 | 5206.2462 | 2603.6267 |
| 739.4171 | --- | 756.4437 | --- | 7 | L | 27 | 5123.1965 | 2562.1019 | 5107.1778 | 2554.0925 |
| 810.4542 | --- | 827.4808 | --- | 8 | A | 26 | 5010.1124 | 2505.5599 | 4994.0937 | 2497.5505 |
| 939.4968 | --- | 956.5234 | --- | 9 | E | 25 | 4939.0753 | 2470.0413 | 4923.0566 | 2462.0319 |
| 1053.5397 | --- | 1070.5663 | --- | 10 | N | 24 | 4810.0327 | 2405.5200 | 4794.0140 | 2397.5106 |
| 1216.6031 | --- | 1233.6296 | --- | 11 | Y | 23 | 4695.9898 | 2348.4985 | 4679.9711 | 2340.4892 |
| 3407.3962 | --- | 3424.4228 | --- | 12 | N(HexNAc4Hex6Fuc2) | 22 | 4532.9265 | 2266.9669 | 4516.9077 | 2258.9575 |
| 3535.4912 | 1768.2492 | 3552.5178 | 1776.7625 | 13 | K | 21 | 2342.1333 | 1171.5703 | 2326.1146 | 1163.5609 |
| 3622.5232 | 1811.7653 | 3639.5498 | 1820.2785 | 14 | S | 20 | 2214.0383 | 1107.5228 | 2198.0196 | 1099.5134 |
| 3737.5502 | 1869.2787 | 3754.5767 | 1877.7920 | 15 | D | 19 | 2127.0063 | 1064.0068 | 2110.9876 | 1055.9974 |
| 3851.5931 | 1926.3002 | 3868.6197 | 1934.8135 | 16 | N | 18 | 2011.9794 | 1006.4933 | 1995.9606 | 998.4840 |
| 4011.6238 | 2006.3155 | 4028.6503 | 2014.8288 | 17 | C(Carbamidomethyl) | 17 | 1897.9364 | 949.4719 | 1881.9177 | 941.4625 |
| 4140.6663 | 2070.8368 | 4157.6929 | 2079.3501 | 18 | E | 16 | 1737.9058 | 869.4565 | 1721.8871 | 861.4472 |
| 4255.6933 | 2128.3503 | 4272.7198 | 2136.8636 | 19 | D | 15 | 1608.8632 | 804.9352 | 1592.8445 | 796.9259 |
| 4356.7410 | 2178.8741 | --- | --- | 20 | T | 14 | 1493.8362 | 747.4218 | 1477.8175 | 739.4124 |
| 4453.7937 | 2227.4005 | 4470.8203 | 2235.9138 | 21 | P | 13 | 1392.7886 | 696.8979 | --- | --- |
| 4582.8363 | 2291.9218 | 4599.8629 | 2300.4351 | 22 | E | 12 | 1295.7358 | 648.3715 | 1279.7171 | 640.3622 |
| 4653.8734 | 2327.4404 | 4670.9000 | 2335.9536 | 23 | A | 11 | 1166.6932 | 583.8502 | 1150.6745 | 575.8409 |
| 4710.8949 | 2355.9511 | 4727.9215 | 2364.4644 | 24 | G | 10 | 1095.6561 | 548.3317 | 1079.6374 | 540.3223 |
| 4873.9582 | 2437.4828 | 4890.9848 | 2445.9960 | 25 | Y | 9 | 1038.6346 | 519.8210 | 1022.6159 | 511.8116 |
| 5021.0266 | 2511.0170 | 5038.0532 | 2519.5302 | 26 | F | 8 | 875.5713 | 438.2893 | 859.5526 | 430.2799 |
| 5092.0638 | 2546.5355 | 5109.0903 | 2555.0488 | 27 | A | 7 | 728.5029 | 364.7551 | 712.4842 | 356.7457 |
| 5205.1478 | 2603.0776 | 5222.1744 | 2611.5908 | 28 | I | 6 | 657.4658 | 329.2365 | 641.4470 | 321.2272 |
| 5276.1849 | 2638.5961 | 5293.2115 | 2647.1094 | 29 | A | 5 | 544.3817 | 272.6945 | 528.3630 | 264.6851 |
| 5375.2534 | 2688.1303 | 5392.2799 | 2696.6436 | 30 | V | 4 | 473.3446 | 237.1759 | 457.3259 | 229.1666 |
| 5474.3218 | 2737.6645 | 5491.3483 | 2746.1778 | 31 | V | 3 | 374.2762 | 187.6417 | 358.2575 | 179.6324 |
| 5602.4167 | 2801.7120 | 5619.4433 | 2810.2253 | 32 | K | 2 | 275.2078 | 138.1075 | 259.1890 | 130.0982 |
| --- | --- | --- | --- | 33 | K | 1 | 147.1128 | 74.0600 | 131.0941 | 66.0507 |

**Figure S8.** Examples of N-glycopeptide localization in Protein Prospector. (A) shows a completely localized GSM; (B) GSM with the correct sequence but incorrect glycan localization, and (C) a GSM with insufficient localization and sequence information. Since Protein Prospector does not consider more than one fragmentation type, only the EThcD spectrum are shown. In (A), c/z fragment ions confirmed both peptide sequence and glycan localization. In (B), the peptide sequence was confirmed using b/y and c/z fragments, but fragments were not detected between the Asn and Ser. Therefore, we could not confidently localize the glycan. Finally, in (C), very little peptide fragmentation corresponding to the proposed peptide was seen in the EThcD spectrum. All examples in this figure correspond to the glycoprotein mixture, and annotated spectra were generated using the viewer available in the Protein Prospector software suite.

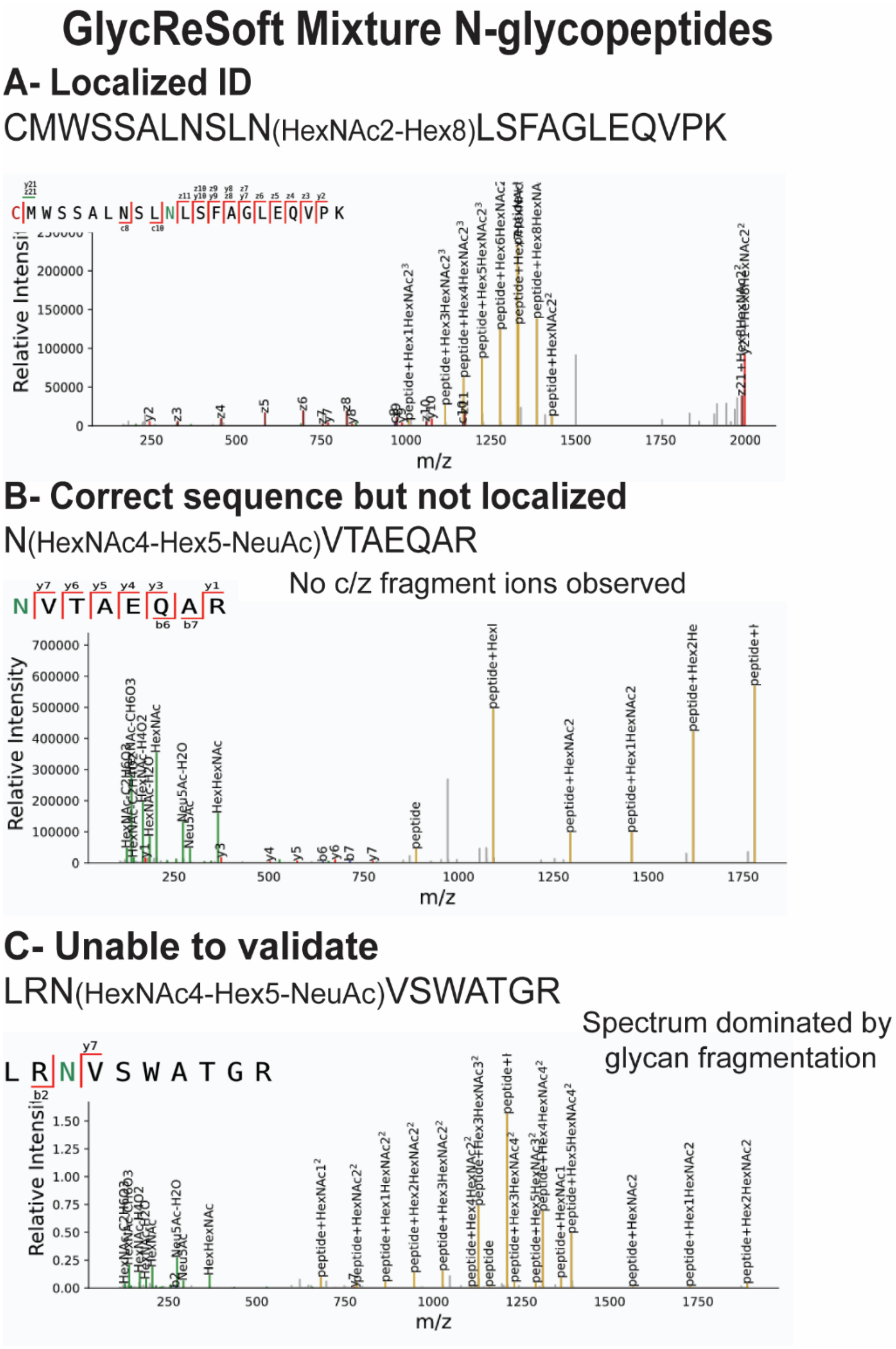


**Figure S9.** Examples of N-glycopeptide localization in GlycReSoft. (A) shows a completely localized GSM; (B) GSM with the correct sequence but incorrect glycan localization, and (C) a GSM with insufficient localization and sequence information. In (A) c/z and b/y fragment ions confirm both peptide sequence and glycan localization. In (B), the peptide sequence was confirmed using b/y fragment ions, but no fragments were observed between the Asn and Thr. Therefore, we could not confidently localize the glycan. Finally, in (C), very little peptide fragmentation corresponding to the proposed peptide was seen in the EThcD spectrum. All examples in this figure correspond to the glycoprotein mixture, and annotated spectra were generated using the viewer available in the GlycReSoft software suite.

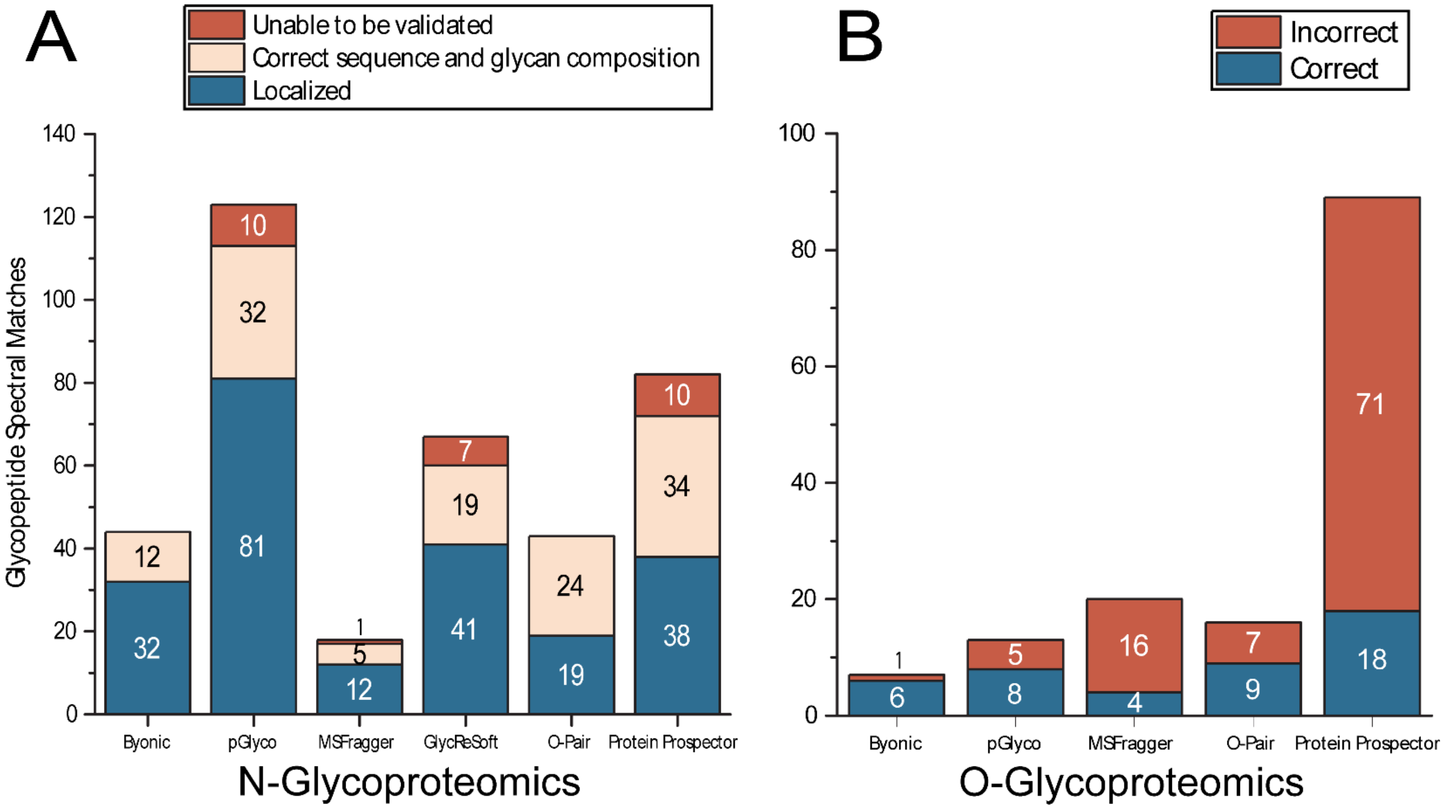


**Figure S10.** Bar graph representing validation of N- (A) and O- (B) GSMs identified by six search algorithms in the complex mixture sample. Filtering parameters are described in the methods section of the main text. In A, blue signifies GSMs that were localized using EThcD, light pink signifies GSMs where the peptide sequence and glycan compositions were validated, and dark red signifies GSMs that were unable to be validated. In B, blue signifies O-GSMs that were localized using EThcD and dark red signifies GSMs that were unable to be localized.

| **Fetuin - P12763** | | **Sites** | **Total** |
|---|---|---|---|
| **Predicted or reported** | N - Uniprot | 99, 156, 176 | **3** |
| | O - NetOGlyC | 253, 271, 280, 282, 290, 296, 314, 216, 320, 323, 324, 325, 334, 341 | **14** |
| **O-Pair** | N- | 156 | **1** |
| **pGlyco** | N- | 156 | **1** |
| **MSFragger** | N- | 156 | **1** |
| **Protein Prospector** | N- | 156 | **1** |
| | O- | 341, 334 | **2** |
| **GlycReSoft** | N- | 156 | **1** |
| **Byonic** | N- | 156 | **1** |

**Table S1.** N- and O-glycosite analysis of fetuin. N-glycosylation sites (3) were collected by Uniprot and O-glycosylation sites (14) were predicted with NetOGlyc[1]. All software (O-Pair, pGlyco, MSFragger, Protein Prospector, GlycReSoft and Byonic) reported 1 N-glycosylation site and Protein Prospector found 2 O-glycosylation sites. Experimental glycosites reported here were manually validated.

| CD14 - P08571 | | Sites | Total |
|---|---|---|---|
| **Predicted or reported** | N - Uniprot | 37, 151, 282, 323 | **4** |
| | O - NetOGlyC | 129, 153, 182, 264, 346, 362 | **6** |
| **O-Pair** | N- | 151 | **1** |
| **pGlyco** | N- | 151, 282 | **2** |
| **MSFragger** | N- | 282 | **1** |
| **Protein Prospector** | N- | 151 | **1** |
| **GlycReSoft** | N- | 151, 282 | **2** |
| **Byonic** | N- | 151, 282 | **2** |

**Table S2.** N- and O-glycosite analysis of CD14. N-glycosylation sites (4) were collected by Uniprot and O-glycosylation sites (6) were predicted using NetOGlyc[1]. A total of 2 N-glycosylation sites were identified; GlycReSoft, Byonic, and pGlyco identified both, while O-Pair, MSFragger and Protein Prospector identified a single N-glycosite. None of the programs reported any O-glycosites. Experimental glycosites here were manually validated.

| Serotransferrin - P02787 | | Sites | Total |
|---|---|---|---|
| **Predicted or reported** | N - Uniprot | 432, 491, 630 | **3** |
| | O - NetOGlyC | 306, 561 | **2** |
| | O - Uniprot | 51 | **1** |
| **O-Pair** | N- | 630 | **1** |
| **pGlyco** | N- | 432 | **1** |
| **MSFragger** | N- | 432, 630 | **2** |
| **Protein Prospector** | N- | 432, 630 | **2** |
| | O- | 24 | **1** |
| **GlycReSoft** | N- | 432, 630 | **2** |
| **Byonic** | N- | 432, 630 | **2** |

**Table S3.** N- and O-glycosite analysis of serotransferrin. N-glycosylation sites (3) were collected by Uniprot and O-glycosylation sites (3) were predicted with NetOGlyc[1]. A total of 2 N- and 1 O-glycosites were identified. MSFragger, Protein Prospector, GlycReSoft and Byonic identified both N-glycosites, while O-Pair and Protein Prospector identified one of the sites. Protein Prospector was the only search algorithm to identify an O-glycosylation site. Experimental glycosites reported here were manually validated.

| ApolipoproteinE - P02649 | | Sites | Total |
|---|---|---|---|
| **Predicted or reported** | O - NetOGlyC | 26, 35, 40, 157, 193, 212, 215, 241, 307, 308 | **10** |
| **O-Pair** | O- | 212, 308 | **2** |
| **pGlyco** | O- | 212, 308 | **2** |
| **MSFragger** | O- | - | **0** |
| **Protein Prospector** | O- | 212, 307, 308 | **3** |
| **Byonic** | O- | 212, 308 | **2** |

**Table S4.** N- and O-glycosite analysis of Apolipoprotein E. O-glycosylation sites (10) were predicted with NetOGlyc[1]. O-Pair, pGlyco and Byonic identified 2 O-glycosylation sites while Protein Prospector identified 3. Experimental glycosites reported here were manually validated.

| Coagulation Factor 12 - P00748 | | Sites | Total |
|---|---|---|---|
| Predicted or reported | N- | 249,433 | **2** |
| | O- | 17, 19, 62, 75, 76, 99, 102, 153, 175, 216, 224, 231, 232, 234, 243, 246, 251, 297, 299, 305, 308, 238, 329, 331, 335, 337, 350, 352, 358, 366, 369 | **27** |
| O-Pair | O- | 327, 328, 335, 337 | **4** |
| | N- | 249,433 | **2** |
| pGlyco | O- | 327, 328, 335, 337 | **4** |
| | N- | 249,433 | **2** |
| MSFragger | O- | 115, 328 | **2** |
| | N- | 249, 433 | **3** |
| GlycReSoft | N- | 249,433 | **2** |
| Protein Prospector | O- | 24, 109, 153, 329, 331, 335, 337 | **7** |
| | N- | 249,433 | **2** |
| Byonic | O- | 327, 328, 337, 335 | **3** |
| | N- | 249,433 | **2** |

**Table S5.** N- and O-glycosite analysis of coagulation factor 12. O-glycosites (27) were predicted using NetOGlyc[1]. O-Pair, pGlyco and Byonic identified 4 O-glycosylation sites, while Protein Prospector identified 7 and MSFragger identified 2. Experimental glycosites reported here were manually validated.

| von Willebrand factor - P04275 | | Sites | Total |
|---|---|---|---|
| Predicted or reported | N- | 90, 156, 211, 666, 857, 1231, 1515, 1574, 2223, 2290, 2357, 2400, 2546, 2585, 2790 | **15** |
| | O- | 204, 207, 208, 209, 214, 292, 306, 314, 362, 567, 577, 580, 588, 589, 600, 668, 671, 741, 754, 755, 758, 761, 764, 766, 1028, 1034 1041, 1042, 1045, 1054, 1119, 1213, 1217, 1248, 1253, 1255, 1256, 1468, 1477, 1486, 1487, 1679, 1901, 1914, 1926, 1930, 1932, 2202, 2225, 2226, 2247, 2292, 2293, 2298, 2303, 2347, 2349, 2359, 2371, 2374, 2382, 2386, 2404, 2412, 2413, 2415, 2421, 2422, 2423, 2424, 2435, 2436, 2469, 2479, 2482, 2503, 2505, 2510, 2512, 2513, 2516, 2519, 2523, 2548, 2559, 2568, 2594, 2602, 2613, 2623, 2660, 2666, 2691, 2775, 2779 | **97** |
| O-Pair | N- | 857, 2223, 2585 | **3** |
| pGlyco | N- | 857, 2790, 2585, 1515, 2357 | **5** |
| MSFragger | O- | 859 | **1** |
| | N- | 2223, 2790, 857 | **3** |
| GlycReSoft | N- | 2223, 857, 2400, 2790 | **4** |
| Protein Prospector | N- | 857, 2223, 2357 | **3** |
| Byonic | N- | 857, 2357, 2223, 2585 | **4** |

**Table S6.** N- and O-glycosite analysis of von Willebrand factor. O-glycosites (97) were predicted using NetOGlyc[1] and N-glycosylation sites (15) were collected by Uniprot. pGlyco identified 5 N-glycosites, followed by GlycReSoft and Byonic, which identified 4 each, and finally, MSFragger and Protein Prospector identified 3 each. MSFragger was the only software to identify an O-glycosite. Experimental glycosites reported here were manually validated.
